# Nonrandom interchromatin trafficking through dynamic multiphase speckle connections

**DOI:** 10.1101/2025.05.23.655761

**Authors:** Jiah Kim, Gabriela Andrea Hernandez Gonzalez, Neha Chivukula Venkata, Kyu Young Han, Andrew S. Belmont

## Abstract

Nuclear speckles (NS) enhance the expression of NS-associated genes, possibly by elevating local levels of factors involved in multiple steps of gene expression. While dozens of large NS are distributed throughout interchromatin regions, the extent to which NS components dynamically redistribute between NS to adjust to local physiological demands remains unknown. Here we used live-cell imaging of endogenous NS proteins to identify an interchromatin network of connections that functionally link NS throughout the nucleus. Over timescales ranging from tens of seconds to minutes, NS material undergoes bulk transfer through these connections. Multiphase NS-connecting structures form through the dynamic juxtapositions of multiple NS component phases. Each phase exhibits distinct yet recurrent viscoelastic dynamics, but together, they integrate into a more stable, multiphase, NS-connecting structure in an ATP-and transcription-dependent manner. Our findings reveal the existence of a cellular mechanism that facilitates coordinated inter-NS protein trafficking through multiphase connections.

## Introduction

The eukaryotic cell nucleus achieves functional compartmentalization through spatial segregation of nuclear components^1-3^. This compartmentalization includes the compaction of chromatin into discrete, locally condensed, large-scale chromatin domains positioned along the interchromatin (IC) space^2,4^. An intriguing aspect of this large-scale chromatin domain-IC space organization is its possible functional role in facilitating the movement of large macromolecules such as mRNPs, which preferentially travel through the interchromatin space^5-10^.

Nuclear speckles (NS) are large nuclear condensates embedded within the IC space^11-13^, associating with active transcription sites and euchromatin regions^14,15^. NS proteomics reveals an enrichment of hundreds of factors involved in different stages of gene expression, especially different steps of RNA processing and nuclear export^16-18^. Multiple orthogonal genomic approaches have revealed specific NS-associated chromosome regions that show elevated overall levels of gene expression and are greatly enriched in the most highly expressed genes^14,19-21^. Similarly, multiple orthogonal genomic methods have identified specific RNAs that either transiently traffic through NS or are conditionally retained in NS^18,22^. Thus, the original proposal that NS act as a functional "hub" enhancing and regulating the expression of NS-associated genes^11^ is now increasingly supported by the genome-wide intranuclear localization mapping of DNA and RNA as well as the experimental demonstration of gene expression amplification^23^ and increased splicing efficiency^24^ of NS-associated genes^15^.

Given this increasing support for NS acting as bona fide gene expression hubs, new questions arise. One of these is how formation of NS within the IC space and trafficking of factors in and out of NS and between NS might be related to changing levels of nearby gene expression. NS behave in many ways as biomolecular condensates and thus show rapid diffusion of component molecules into and out of NS ^25-30^. Other behaviors of NS include fusion, fission, and shape changes^31,32^, including protrusions that may extend toward active gene^33^. More specifically, inhibition of transcription induced increased movement of NS through the IC space and fusion with neighboring, larger NS. These NS movements repeatedly occurred along similar paths within the IC space, marked by elevated numbers of SON protein "granules"^31^. This observation suggested the possibility that, beyond the observed movement of NS, there might also be movement of bulk NS material between NS within specific IC channels. However, exactly how NS might be spatially connected and whether such connectivity might facilitate non-diffusive, directional transfer of components between neighboring NS remained unknown.

Here we used a combination of wide-field (WF) live cell microscopy, photoactivation, and single-molecule tracking (SMT) to visualize the directional spreading of NS materials specifically through subregions of the IC space lying between adjacent NS. The persistent maintenance and repeated formation of these NS connections under normal cell growth conditions occurs through the ATP and transcription dependent spatial juxtapositions of multiple condensate phases. These multiple condensate phases dynamically rearrange relative to each other while maintaining a linear bridge spanning adjacent NS that appear to support transport of NS materials over micron-scale distances and over a tens-of-seconds to minutes timescale. We speculate that these multiphase connections link individual NS into a larger functional network allowing for dynamic reallocation of gene expression factors throughout the nucleus to adjust to local variations in gene expression.

## RESULTS

### Visualization of SON-enriched connections between NS

We used CRISPR-Cas9 to insert EGFP or mEOS3.2 fluorescent protein sequences into the N-terminus coding sequence of the endogenous SON gene. SON and SRRM2 are core NS components not only because of their high NS enrichment, and interior localization^16,34^, but also due to their essential role in NS formation; simultaneous knockdown (KD) of SON and SRRM2 disperses multiple other NS proteins, suggesting loss of NS after KD^35^.

By live cell WF microscopy (**Fig. 1a**, left panel), the GFP-tagged endogenous SON protein appears ∼5-10-fold more concentrated within NS relative to its mean nucleoplasmic level. This ratio is an underestimate, though, due to the limited z-resolution of WF deconvolution microscopy. Lower levels of SON protein outside of NS are not uniformly distributed but rather concentrate within DNA-depleted regions, particularly between neighboring NS. Due to the nonlinearity of human perception of intensity differences, we generated a pseudo-color intensity map to more clearly visualize low-intensity SON accumulations outside of NS. We used 10 discrete intensity levels, designating any intensity exceeding 30% of the maximum intensity in the field of view as 10 (**Fig. 1a**, right panel) with the remaining intensity range divided into 9 additional levels, with the SON nuclear background intensity falling within level 1. This approach highlights subtle SON signals, including small condensates and foci, revealing a nonrandom distribution of lower intensity SON signals spanning roughly linear paths between particular adjacent NS.

**Figure 1.**
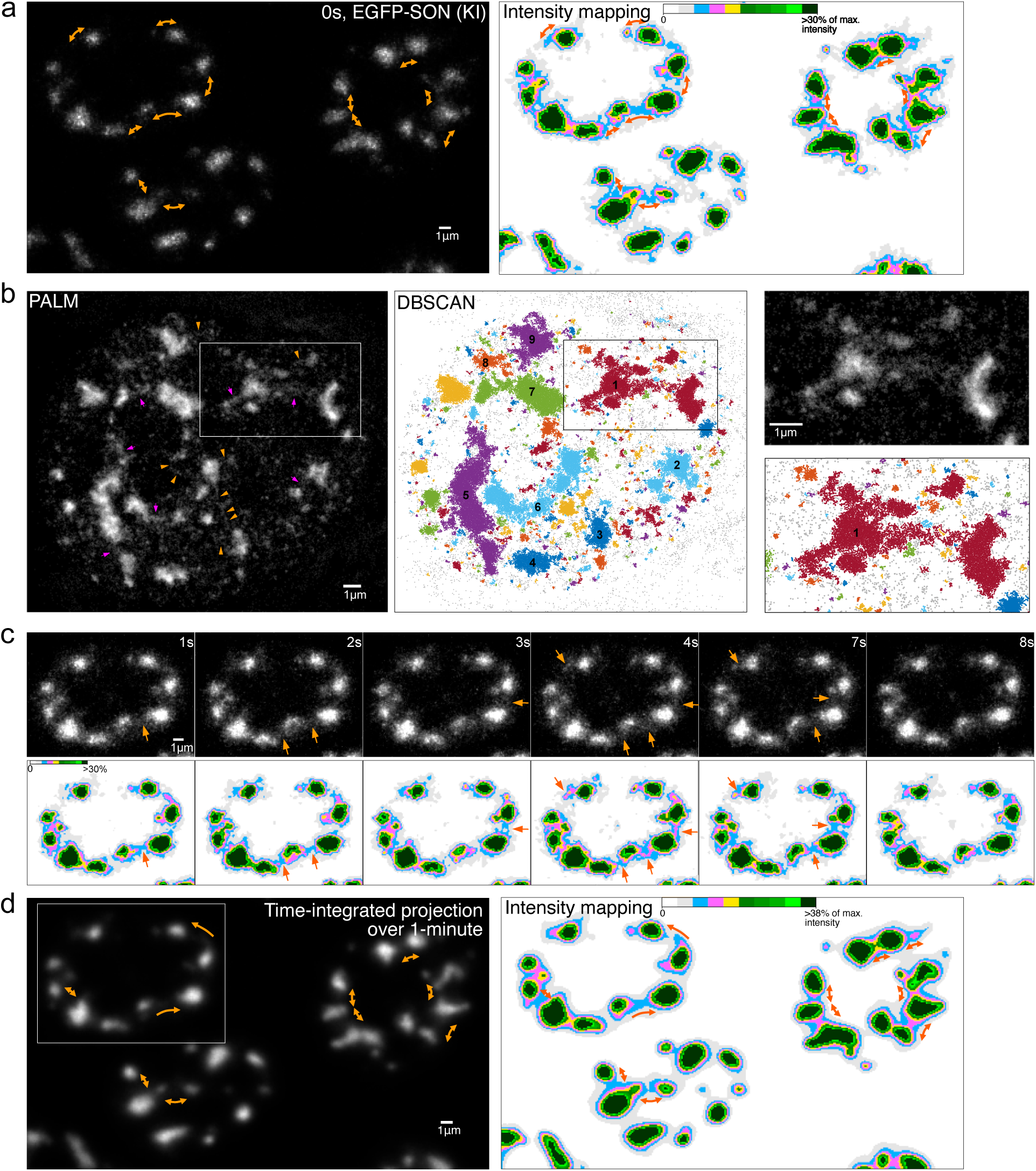
Live-cell wide-field and super-resolution light microscopy reveals NS connections. **a.** Wide-field (WF) deconvolution microscopy of endogenous EGFP-SON in live-cell (**Video 1**). Left: 1^st^ time-lapse image of time series. Right: 10-level pseudo-color intensity mapping of same image. Arrows show where protrusions of transiently elevated SON intensities appear between NS (blue) within broader regions with lower SON intensities (gray). **b.** 2D PALM images, segmented into color-coded clusters of detected single molecules (right), reveal grouping of multiple NS connected by lower-density SON linkages. Left: PALM image of mEOS3.2-SON, expressed from the endogenous gene, reconstructed from 10-min live-cell acquisition. Orange arrowheads point to linear tracks of SON molecules spanning adjacent NS. Magenta arrowheads point to broader linear zones of SON molecules, also spanning NS. Middle: DBSCAN cluster analysis of the PALM image (k=5, ε=50 nm; see SFig.1 for other parameters). Color-coded clusters reveal multiple NS connected by lower-density SON distributions. Right: Higher-magnification view of the boxed region in left and middle panels, showing the original PALM image (top) and DBSCAN-identified clusters (bottom). **c.** Time-lapse images of EGFP-SON showing formation of transient NS connections. Top: non-linear grayscale images with *γ* =0.5 effectively reduce the intensity dynamic range, enhancing visibility of low intensity signals. Bottom: 10-level pseudo-color intensity mapping of linearly scaled images. Arrows point to lower intensity SON regions typically bridging adjacent NS. **d.** Time-integrated projection (sum-intensity) of 1-min live cell images (60 frames, left) and corresponding intensity mapping (right) reveal varying connectivity between different NS. The boxed nucleus is the same as in (**c**), and SFig. 2. Bi-directional arrows point to established NS connections, while single-directional arrows show direction of emerging connections.

To better visualize this lower intensity SON distribution in live cells, we used Photoactivated Localization Microscopy (PALM) (**Fig. 1b**). In these higher-resolution PALM images, acquired over a 10-minute period, we observed concentrations of SON molecules within NS; additional SON molecules aligned into local linear accumulations that frequently extended outward from these NS and formed bridges between neighboring NS and SON foci across micron-scale distances (**Fig. 1b**, orange arrowheads). In some regions, these accumulations appeared to merge, forming broader, higher-density structures aligned between neighboring NS. (**Fig. 1b**, magenta arrows).

To more objectively visualize apparent tracks of SON molecules between NS from these PALM point distributions, we applied the Density-Based Spatial Clustering of Applications with Noise (DBSCAN) algorithm that groups SON localization by density^36^. Each cluster was defined as the set of points with at least 5 neighboring points (k) within a 50 nm radius (ε) (**Fig. 1b**, middle panel). Using these parameters, many clusters corresponded to small NS and/or their protrusions, but some spanned multiple NS suggesting NS connectivity. (**Fig. 1b**, i.e., clusters 1, 2, 5, 6 and 7 in middle and right panels). When these DBSCAN parameters were relaxed, neighboring clusters defined by the original parameters merged, connecting more neighboring NS within local nuclear regions into single clusters (**SFig. 1**).

We next examined the dynamics of these SON densities between NS using time-lapse WF live-cell imaging. We observed SON accumulations between neighboring NS repeatedly reforming over a timescale of seconds (**Fig. 1c, Video 1**).

Pseudo-color intensity maps showed that higher-intensity connections (blue-pink-yellow) formed transient, thinner protrusions (**Fig. 1c**, lower panels), whereas lower-intensity SON accumulations (gray) surrounding and extending between NS remained stable. These transient connections frequently appeared at various inter-NS locations, disappearing and then reappearing at the same sites over time, while remaining within the broader, stable, lower-intensity bridge (**Fig. 1c**, lower panels, orange arrows, **Video 1**).

To better appreciate this recurrence of connections and protrusions over the same regions between neighboring NS, we generated time-integrated projections by summing individual time-lapse images over multiple time points (**Fig. 1d, SFig. 2**). A 1-minute time-integrated projection (1 frame/sec) showed more continuous and numerous connections between neighboring NS with elevated SON-intensity compared to individual time-point images. These projections highlighted regions where connections formed more frequently. In some regions, they revealed directional bridge formation from one NS to another (**Fig. 1d**, one-directional arrow, **SFig. 2, Video 2),** whereas, in other regions, bridges formed through connections emanating from both NS (**Fig. 1d**, bidirectional arrows).

**Figure 2.**
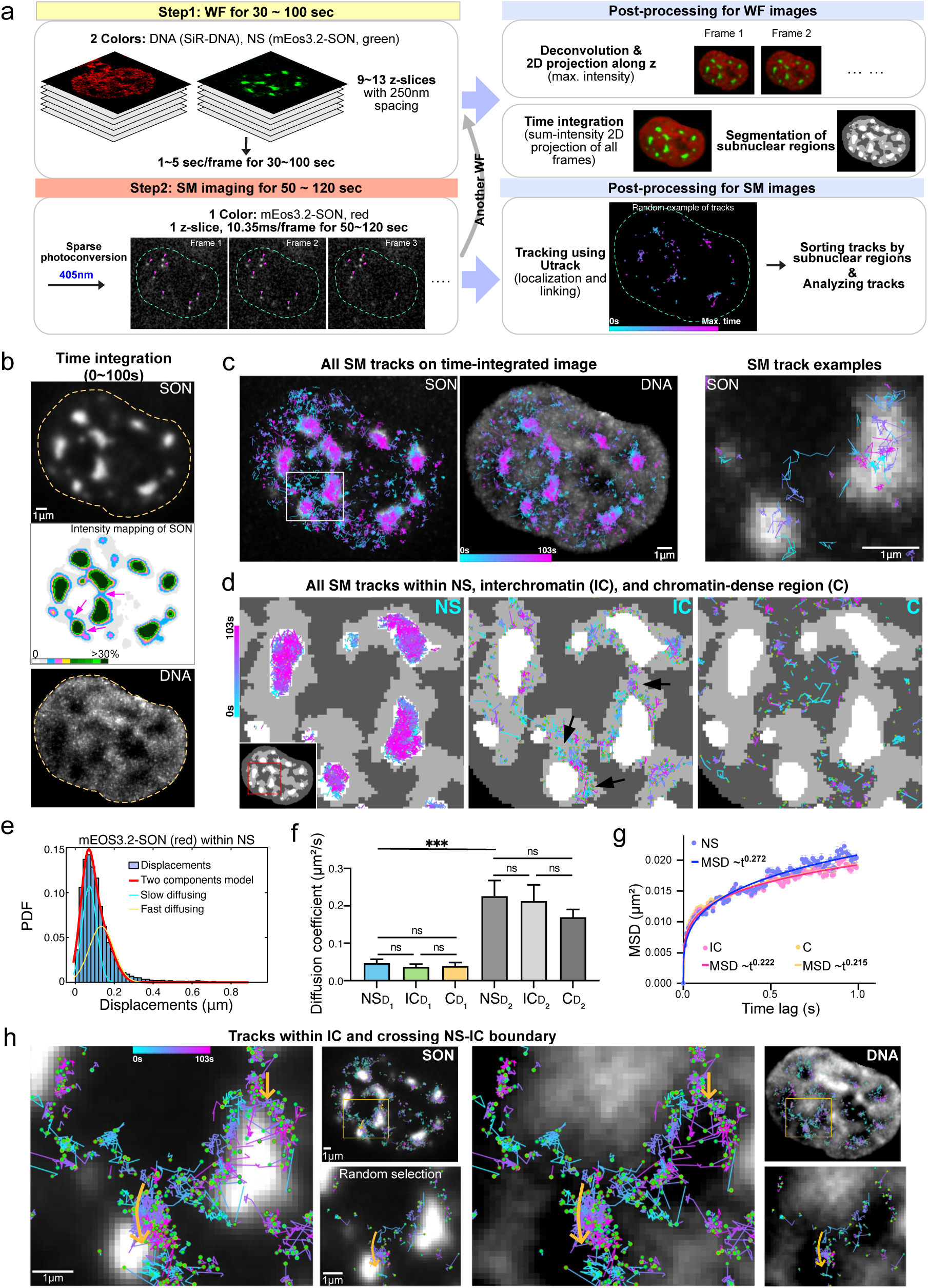
Both NS and NS-connecting regions show similar SON molecular movements. **a.** Schematic illustrating imaging and post-processing steps for combining Single Molecule Tracks (SMT) with wide-field (WF) visualization of nuclear structure. **b.** Time-integrated projection (sum-intensity) over 100-secs identifies NS, NS-connecting regions, and DNA-dense regions. Top: time-integrated projection of mEos3.2-SON over 100s; Middle: 10-level pseudo-color mapping of the same projection. Arrows indicate NS connections. Bottom: Time-integrated projection of DNA stained with SiR-DNA. Dotted lines outline nucleus. **c.** SMT color-coded by occurrence time, superimposed on the time-integrated projection of SON (left) and DNA (middle). Enlarged view (right) showing randomly selected SMT superimposed on time-integrated projection of SON from the boxed nuclear region (left)**. d.** Sorting of SMT contained solely within NS, interchromatin (IC), or chromatin-dense regions (C). Left, middle, right panels show SMT (colored) superimposed over segmented NS (white), IC (light gray), and C (dark gray) regions, respectively, from boxed region in lower-magnification view of entire nucleus (inset, left panel). Black arrows (middle panel) point to regions shown in (**b**) where NS connections more frequently form. **e.** Distribution of frame-to-frame displacements for SMT confined within NS. Fitting of distribution used a two-diffusing state model: overall population-red line (R^2^=0.989, RMSE=0.0043), slow subpopulation-cyan line, faster subpopulation-yellow line. **f.** Estimated diffusion coefficients for slow-moving (D_1_) and fast-moving (D_2_) subpopulations are similar for SON molecules within NS, IC, and C regions. N = 16 cells (44788 tracks in NS, 18852 in IC, 17810 in C). One-way ANOVA Kruskal-Wallis test (ns p>0.05, *** p<0.001) **g.** Mean square displacement (MSD) plots versus time of SON (Halotag-JFX554) SMT from different nuclear regions with fitting (R^2^=0.999, solid lines) to MSD=4Dt^α^. (N = 18 cells, 9377 NS tracks, 18310 IC tracks, 21656 C tracks). **h.** SMT within IC, including those crossing NS-IC boundary, superimposed on the SON (left panels) or DNA (right panels) time-integrated projections from a nuclear subregion (yellow boxes, upper right image of each panel). The lower right image of each panel shows randomly selected SMT subpopulation. Green dots show where the SMT starts. Yellow arrows indicate groups of SMT showing apparent progressive SON translocation towards NS as indicated by color shifts from early (cyan, blue) to later (purple, pink) times points.

In summary, by combining WF and PALM live-cell imaging we identified transient elevations of SON intensity that bridge neighboring NS. In contrast to conventional nuclear bodies, these SON-containing connections are short-lived, but show a unique form of stability - repeatedly forming at the same or nearby location, as appreciated in time-averaged projections.

### The molecular movements of SON within interchromatin regions suggest a gradual bulk transfer of SON between NS

The elevated SON intensities in transient connections between NS raised questions about the molecular movements of SON within these regions. First, how does SON mobility within these transient connections compare to that within the NS interior, the IC space, or chromatin dense regions? Second, might these transiently elevated SON intensities reflect correlated movements of multiple SON molecules between NS, possibly through bulk flow of NS material.

To compare SON mobility within different nuclear regions, we first identified these different regions by performing two-color WF live-cell imaging of mEOS3.2 tagged endogenous SON (green state) and SiR-DNA staining for 30–100 seconds before and after single-molecule (SM) imaging (**Fig.2a**). As in **Fig. 1d**, 100-second time-projections of the WF live-cell images highlighted nuclear regions in which specific NS were more frequently connected (**Fig. 2b**, arrows, **SFig. 3a**, individual time-lapse images). For SM imaging, inclined illumination^37,38^ was used to image photo-converted mEOS2.3-SON molecules (red state) continuously for 50–120 seconds (**Fig. 2a**).

**Figure 3.**
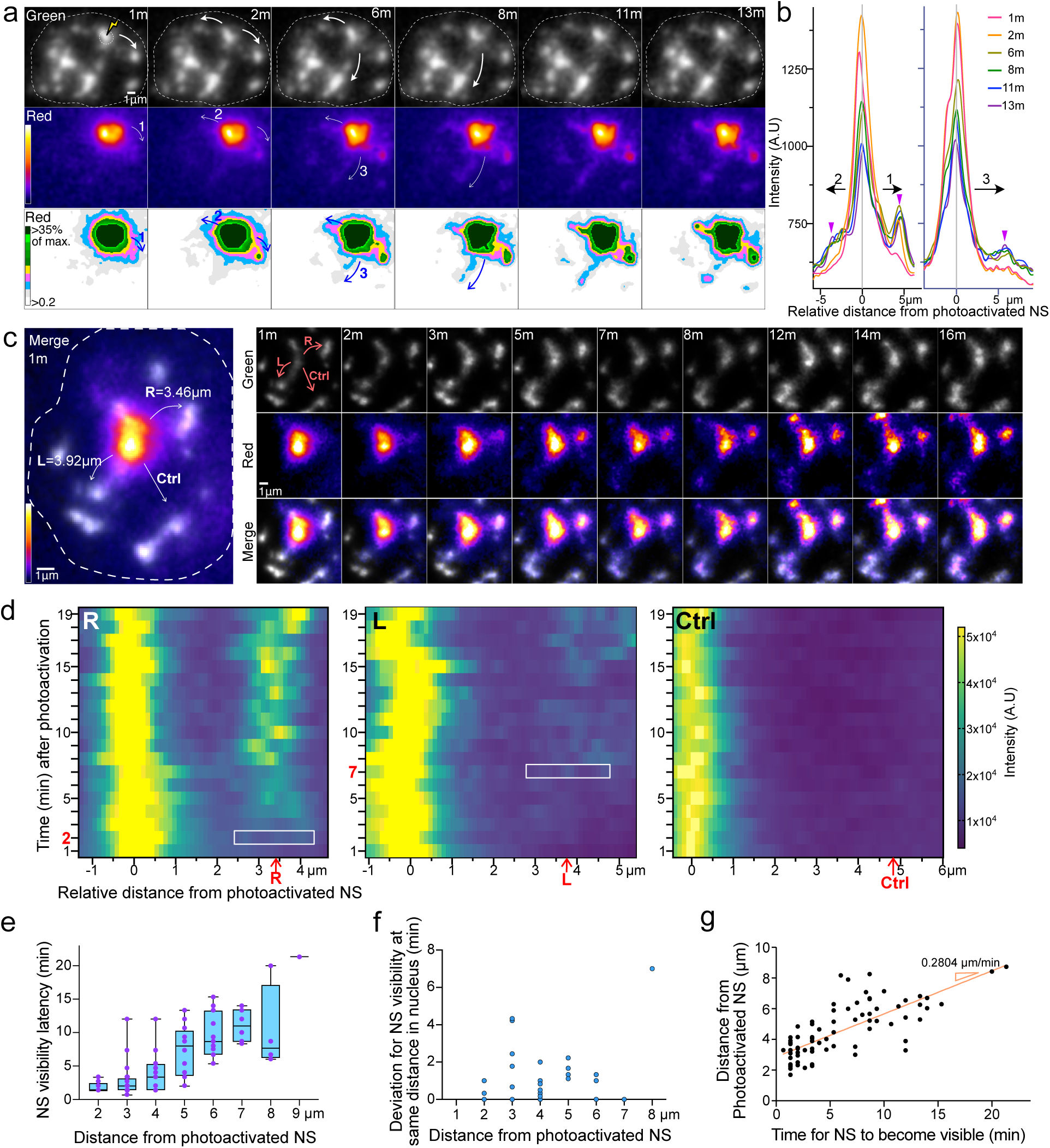
Bulk, directional spreading of SON from photoactivated NS to neighboring NS through NS connections. **a.** Time-lapse images as function of time (m=mins) after photoactivation of one NS (marked by lightning bolt (top-left) show slow, progressive spreading of SON from photoactivated NS to neighboring NS within curvilinear tracks corresponding to locations of low-intensity SON connections between NS identified by imaging of non-photoconverted SON. Top row: non-photoconverted mEos3.2 SON (green) showing overall SON distribution, including bright NS and lower intensity regions-connecting NS. Dotted lines indicate nuclear outline. Middle and bottom rows: photo-converted mEOS3.2-SON (red) displayed as a pseudo-color continuous intensity image (middle) versus a 10-level discrete intensity map (bottom). Images represent sum-intensity z-projections scaled nonlinearly (*γ* = 0.6) to accentuate the low intensity SON connections between NS. Numbered arrows indicate direction and timing order (1-3) of photoactivated SON spreading between different neighboring NS. **b.** Photoconverted SON Intensity line-scans showing SON redistribution over time (different colored lines) along straight lines from the photoactivated NS to the target NS (purple arrowheads). Numbers refer to labeled NS in (**a**). **c.** Variable SON spreading from photoactivated NS (center) to three neighboring NS. Spreading occurs earlier towards "L" than "R" NS, despite equal distance to photoactivated NS; no spreading observed to "Ctrl" NS. Left: merged image at 1 min after photoactivation showing non-photoconverted SON (gray) merged with photoconverted SON (pseudo-colored); Right: time series from 1-16min showing non-photoconverted SON (top), photoconverted SON (middle) and merged images (bottom). All images are sum-intensity z-projection, nonlinearly scaled (*γ* = 0.7) **d.** Pseudo-colored intensity kymograms of photoconverted mEOS3.2-SON line-scan intensities, measured from the photoactivated NS to the target NS (L, R, and Ctrl shown in (**c**)). Photoconverted mEOS3.2-SON signals emerged above background at NS ‘R’ at 2 min and ‘L’ at 7 min (y-axis, time after photoactivation). White boxes mark regions within ±1μm of each target NS, with red arrows and labels indicating NS position (x-axis, distance). No spreading of photoconverted SON above background levels was observed towards NS ‘Ctrl’. **e-f.** Timing of photoconverted mEOS3.2-SON spreading to NS positioned at similar distances is highly variable (statistical comparisons). (**e**) Time required for the target NS to become detectable above background via photoconverted mEOS3.2-SON transfer (y-axis), as a function of distance (x-axis) from the photoactivated NS (N=81 NS from 16 cells). (**f**) Deviation (mins, y-axis) from mean time of photoconverted SON appearance at target NS equidistant within the same nucleus, plotted against distance to photoactivated NS (x-axis, rounded). **g.** Speed of photoconverted SON spreading. Linear regression (red line, R²=0.56) yields a slope of 0.2804 μm/s (data from (**e**)).

SM tracks (SMT) were color-coded by their appearance time during data acquisition and superimposed on time-integrated projections of SON and SiR-Hoechst (DNA) (**Fig. 2c**, **SFig. 3b**). SMT mapped to different nuclear regions proportionally to their SON concentrations, with most SMT occurring within NS. SMT confined entirely within specific segmented nuclear regions-NS, IC space (IC), or chromatin-dense (C) regions-were assigned to their corresponding type regions. (**Fig. 2d**, **SFig. 3b**)

To characterize SON mobility, we calculated apparent diffusion coefficients by fitting the distribution of frame-to-frame displacements from SMT using a two-component model that differentiated between slow and relatively faster diffusing populations of SON molecules (**Fig. 2e**). Both SON mobility and the fraction of slow-versus fast-diffusing populations were similar within all three nuclear regions (**Fig. 2f, SFig. 3c**). These similarities suggest that SON tracks located in different nuclear regions reside within condensates possessing similar physical properties as NS-associated SON condensates. Mean square displacement (MSD) analysis further supported this suggestion. SMT from all three nuclear regions showed similar diffusion exponents ranging between ∼0.22-0.27, indicative of anomalous diffusion (**Fig. 2g**). We therefore conclude that SMT lying within IC spaces are mostly contained within NS-like condensates within the extensions and/or connections between NS visualized by the pseudo-color mapping of WF images (**Fig. 1a**, **Fig. 2b**) and super-resolution PALM images (**Fig. 1b**). Similarly, SMT lying within chromatin dense regions are mostly contained within small SON condensates.

Notably, SMT in IC regions largely localized to areas where NS connections repeatedly formed (**Fig. 2b**, middle panel). Superimposing SMT that spanned both NS and IC regions together with SMT contained entirely within the IC space revealed a network of SMT linking adjacent NS in the time-integrated projection (**Fig. 2h, SFig. 3d**). Within this network we color-coded SMT according to their acquisition time-cyan (early), purple (middle), magenta (late) (**Fig. 2h**).

The difficulty in tracking SON movement continuously from one NS to another over seconds to tens of seconds using SMT is that each SON molecule can typically be tracked for only ∼130ms (13 frames) before photobleaching or leaving the optical section. Nevertheless, we interpreted the temporal and spatial correlation among groups of SON SMT as evidence of progressive SON translocation from one NS to another (**Fig. 2h**), likely occurring within paths corresponding to the low-intensity connections between NS visualized by WF imaging. Within these groups of SMT, individual tracks showed asynchronous, short-lived movements in varying directions as expected for molecular diffusion within liquid-like condensates. Thus, by color-coding SMTs according to their acquisition time, we visualized rapid single-molecule diffusion within condensates, likely translocating slowly along the apparent NS connections outlined by the SMT network.

In summary, we propose that SON transfer between NS on a timescale of seconds to minutes occurs via translocation of SON condensates within which individual SON molecules randomly diffuse.

### Photoactivation experiment demonstrates directional bulk transfer of NS material between individual NS

A priori, the elevated SON intensities connecting NS suggested two possibilities. First, they could reflect the random diffusion of individual SON molecules through the nucleoplasm combined with their transient binding to some other structure or condensate forming connections between NS. Second, they could indicate bulk SON trafficking between NS, occurring through directional translocation of SON condensates between NS, as proposed in the previous section.

To directly test for SON transfer through connections between NS, we performed photoactivation experiments in HCT116 cells expressing mEOS3.2-SON from the endogenous SON locus. Time-lapse imaging tracked the constitutive green fluorescence of mEOS3.2-SON showing the overall nuclear SON distribution, and the red photoconverted fluorescence showing SON spreading from a single photoactivated NS.

After photoactivating a single, large NS, the photoconverted mEOS3.2-SON did not spread rapidly and uniformly from the photoactivated NS, contradicting the random diffusion model. Instead, the photoconverted mEOS3.2-SON formed protrusions that extended progressively over a timescale of several to tens of minutes from the photoactivated NS toward other NS (**Fig. 3a**), as more clearly visualized in the pseudo-color (middle panel) and discrete intensity (bottom panel) images (**Fig. 3a**, **Video 3**). Moreover, these protrusions formed and extended specifically within the pre-existing, low-intensity mEOS3.2-SON connections visualized in the green channel (**Fig. 3a**, top panel).

Furthermore, different protrusions formed at different times, even when connecting to NS approximately equidistant from the photoactivated NS. For example, in **Fig 3a**, photoconverted mEOS3.2-SON spread ∼4-minutes earlier to the NS on the right (NS ‘1’, 2 min) than to the NS on the left (NS ‘2’, 6 min). An even slower, more gradual spreading was observed towards a more distant NS (NS ‘3’). Intensity line profiles further quantified these differences in timing (**Fig. 3b**).

An even greater variation in timing of the photoconverted mEOS3.2-SON spreading to neighboring NS was observed in a second nucleus, with an ∼5-minute difference in the timing of SON spreading to two neighboring NS at comparable distances from the photoactivated NS (**Fig. 3c-d**, ‘R’ at 2 min versus ‘L’ NS at 7 min). In contrast, no spreading occurred towards a third NS (‘Ctrl’) that lacked a visible connection in the green channel showing the overall SON distribution (**Fig. 3c-d**).

We quantified timing delays from photoconversion experiments across multiple nuclei, photoactivating a single NS in each case and measuring the time (mins) required for photoconverted SON signal in each neighboring NS to rise above the background (**Fig. 3e**). Consistent with the two earlier examples (**Fig. 3a-d**), NS at similar distances from the photoactivated NS showed highly variable SON transfer times, with deviations within the same nucleus as high as 7 mins from the mean SON spreading time calculated for all NS within the same distance range (**Fig. 3e-f**). An average photoconverted SON transfer speed of 0.28 μm/min was estimated by the slope of a least-squares fit of NS distance versus transfer time (**Fig. 3g**).

This slow, progressive, yet variably timed protrusion of photoconverted SON within pre-existing NS connections supports the second model of SON bulk trafficking between NS while contradicting the first model for SON trafficking between NS through a diffusive process. More specifically, the model 1 proposal of random diffusion through the nucleoplasm followed by binding to an unknown structure is inconsistent with the observed variable timing of photoconverted SON transfer to neighboring NS at similar distance from the photoactivated NS, as well as the variable timing of photoactivated SON appearance in the connections to these same neighboring NS.

Measurements of the variance in spreading time versus distance were initially based on straight-line distances between NS rather than the contour lengths of the observed curvilinear connections between NS, which correspond to regions of elevated SON concentration. Indeed, the spreading rate of photoconverted SON correlated more closely with curvilinear distances along these pre-existing connections rather than the straight-line distances between NS (**SFig. 4**).

**Figure 4.**
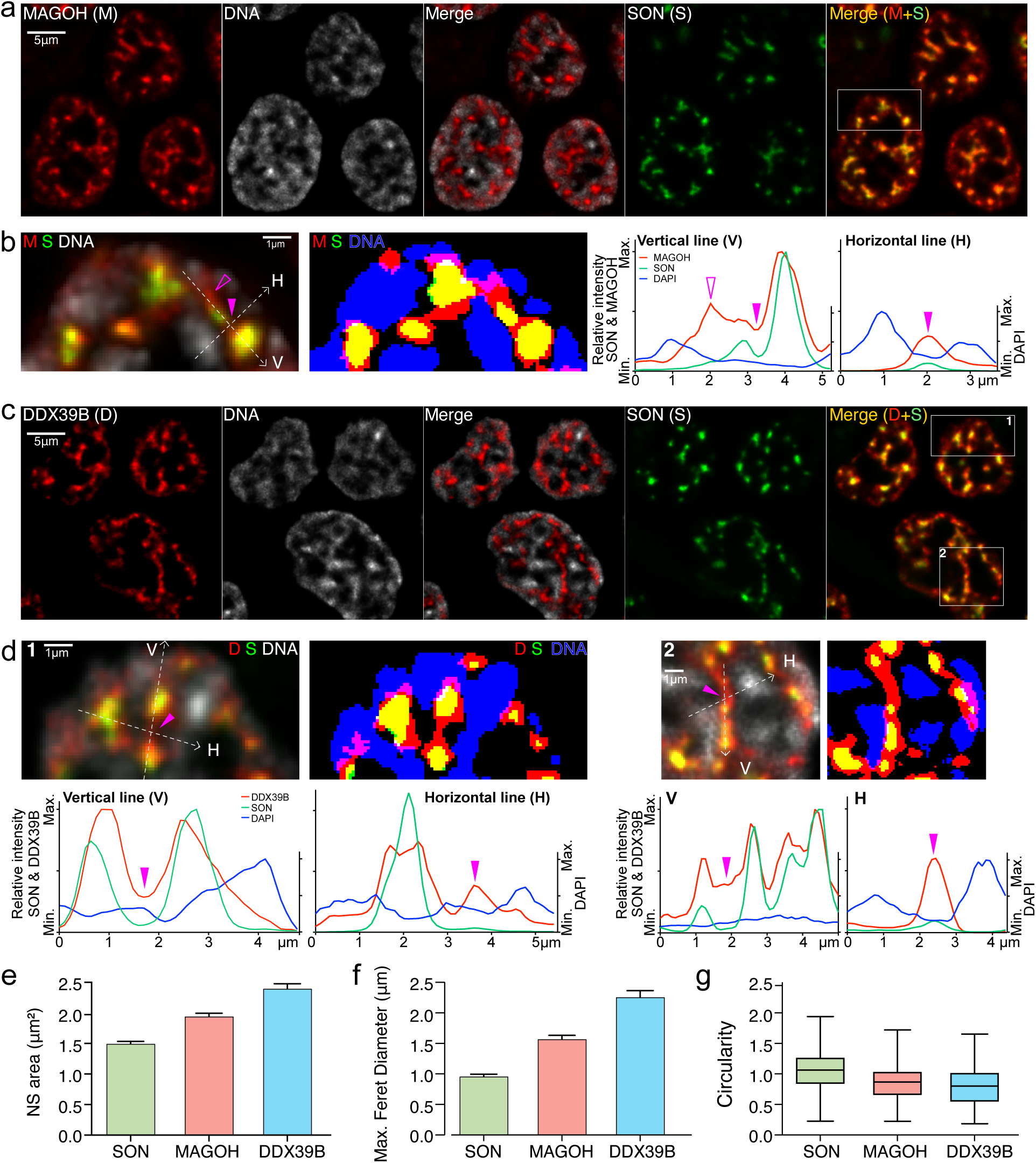
SON speckles are embedded within and connected by MAGOH and DDX39B protein concentrations. **a**&**c.** Representative wide-field images showing Halotag-tagged MAGOH (M, red) (**a**) or DDX39B (D, red) (**c**), EGFP-SON (S, green), all expressed from the endogenous genes, DNA stained with DAPI (grayscale), plus merged images of MAGOH or DDX39B with SON or DNA. **b&d.** Enlarged images and line-scans from boxed nuclear regions from M+S merge image (**a&c**). Merged Images with continuous intensity range (left) are shown next to merged binary images for each color (right). Horizontal (H) and vertical (V) dotted lines indicate the intensity line-scans, either parallel (V) or perpendicular (H) to NS connections. Both MAGOH (**b**) and DDX39B (**d**) show peak intensities within SON speckles, but also elevated levels in connections between NS (solid arrowheads). Relative to their NS intensities, SON intensity is also elevated in these connections, but less than MAGOH and DDX39B. Binary images show that NS proteins form a separate phase within DNA-depleted regions (DAPI). **e-g**. Quantitative comparison of morphologies for the different NS proteins, SON, MAGOH, and DDX39B. Boxplots: mean ± s.e.m. (N= 1251 SON speckles in 155 cells, 780 MAGOH speckles in 87 cells, 573 DDX39B speckles in 68 cells).

In summary, our photoactivation results support bulk flow of SON between NS occurring within specific, curvilinear paths between NS.

### MAGOH and DDX39B overlap with SON-defined NS but additionally surround and bridge between NS

The repeated formation of connections between neighboring NS and the evidence of bulk SON transfer suggested the presence of preferred trafficking pathways. These pathways might be organized by other proteins and/or RNAs, concentrated within the interchromatin space. A significant number of proteins and RNAs have been identified as NS markers ^16,17^, but their abundance and spatial distribution in and around NS vary^34^.

In a concurrent study^39^, we screened for candidate proteins that extended beyond the core SON and SRRM2 NS markers, guided by a proteomics analysis of NS^16^. DDX39B and MAGOH were identified in this screen.

Merged images confirmed that MAGOH and DDX39B indeed formed larger NS that typically colocalized with and surrounded SON but also formed extensions from the NS which often connected NS within the IC space (**Fig. 4, SFig. 5**). Notably, these bridging connections did not simply fill the IC space between NS but corresponded to curvilinear local concentrations of protein of varying width and narrower and more restricted spatially than the total IC space between NS (**Fig. 4a-d**). To compare quantitatively the localization of DDX39B and MAGOH with SON within the IC space, we calculated the Pearson correlation coefficient between various nuclear targets and DAPI-stained DNA (**SFig. 5b**). SON and SRRM2 both negatively correlated with DAPI, demonstrating their predominant localization within DAPI-poor regions. In contrast, histone markers showed higher correlations (>0.5) with DAPI. MAGOH and DDX39B had higher correlation values with DAPI compared to SON but lower correlation values with DAPI than the transcriptionally paused RNA pol2 (RNA pol2S5p), active chromatin (H3K9ac), or facultative heterochromatin (H3K27me3) markers. Both Poly(A) and MALAT1 RNA FISH signals had correlation values with DAPI similar to those of MAGOH and DDX39B, suggesting similar enrichment within the IC space (**SFig. 5**).

**Figure 5.**
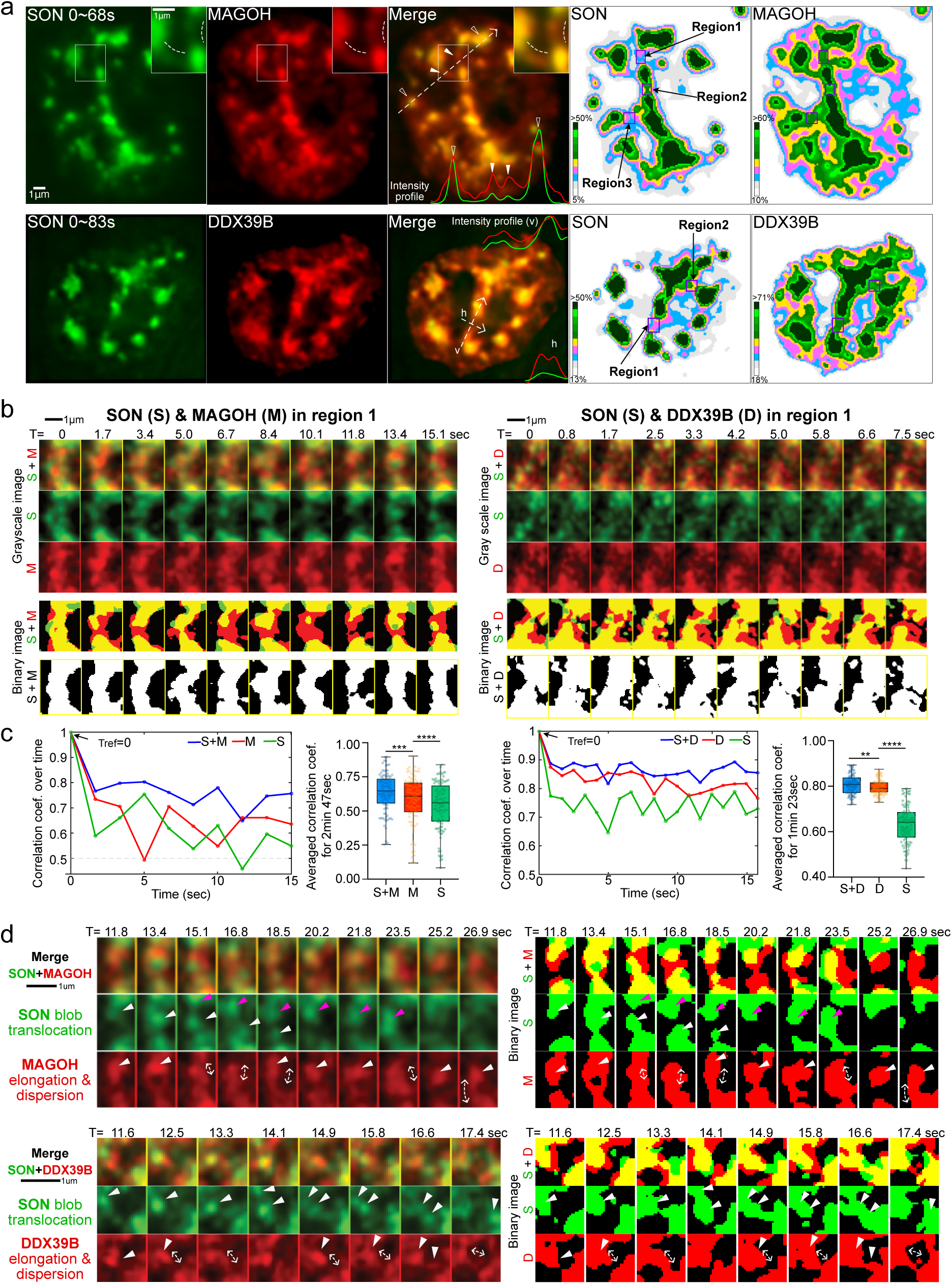
The composite multiprotein phase defines more stable NS connections. **a.** Left panels: Live-cell visualization of NS connections formed by NS proteins SON (left, green), MAGOH (top), and DDX39B (bottom) (middle, red) and merged images (right) using time-integrated projections (sum-intensity; 68 secs for MAGOH, 83 sec for DDX39B). Arrows with white dotted lines correspond to intensity line-scans shown as colored lines in merged image panels. White arrowheads point to NS connections, while non-solid white arrowheads point to NS (top row). Dotted lines in the subpanels (enlarged views of the boxed region in top panels) run parallel to a NS connection. Right panels: 10-level pseudo-color intensity mapping of SON versus MAGOH (top) and SON versus DDX39B (bottom). Boxed regions show NS connection regions of interest analyzed in (**b**) and SFig. 6. **b.** The union of SON and either MAGOH or DDX39B protein phases across the NS connections remains more invariant over time than the distributions of individual proteins. Time-lapse images of individual NS connections, from right panels in (**a**), show merged images (top row), individual proteins (middle rows). Merged binary images show SON (green) versus either MAGOH (left, red) or DDX39B (right, red) segmented regions (2nd from bottom), and their union (bottom row, white). **c.** Spatial correlation analysis of NS connections shown in (**b**). Left panels: Pearson correlation coefficients (left) for SON, MAGOH and their composite signals over time (secs) in region 1 (**b**), using the first frame (T= 0s) as the reference image. The dotted line indicates a correlation coefficient of 0.5. Box plots (right) of averaged cross-correlation coefficients over the 2 min 47 sec imaging period for region 1 (see SFig. 6 for additional regions). Higher and more constant correlation values were observed for the composite NS connection (S+M). Wilcoxon signed-rank test; **p<0.01,***p<0.001,****p<0.0001. Right panels: Same format for SON and DDX39B. (See SFig. 6 for analysis of additional regions). **d.** Examples of different dynamics of individual proteins within NS connection. Arrowheads point to condensate blobs of target proteins, with progressive movement of SON blobs suggesting translocation. Arrows point to apparent elongation or dispersion of a condensate blob visualized in the preceding time frame.

Morphological analysis of segmented images revealed a progressive increase in area and maximum Feret diameters and decreasing circularity for SON, MAGOH, and DDX39B images (**Fig. 4e-g**), consistent with the visualized overlapping but larger coverage of the MAGOH and DDX39B protein distributions which surrounded and sometimes connected adjacent NS (**Fig. 4a-d**).

### Highly dynamic condensates interact to form more stable multiphase NS connections

We next used simultaneous WF imaging of EGFP-SON and either Halotag-MAGOH or Halotag-DDX39B, expressed from their endogenous gene loci in HCT116 cells, to examine the spatial and temporal correlation between SON and MAGOH or SON and DDX39B within NS connections (**Video 4&5**). To better visualize nuclear regions with frequent NS connection formation, we generated time-integrated projections for each color channel over ∼60-90 seconds (**Fig. 5a**, left panels), and then transformed these projections into intensity colormaps (**Fig. 5a**, right panels). In these intensity colormaps, bridges connecting adjacent NS were identified by elevated MAGOH levels (Regions 1–3) or DDX39B levels (Regions 1–2) in green or dark green, with SON present at lower but locally elevated levels, ranging from blue to pink. At individual time points, the spatial patterns of SON and MAGOH in these regions were highly dynamic (**Fig. 5b**, left, **SFig. 6a**). The same was true for the spatial patterns of SON and DDX39B (**Fig. 5b**, right, **SFig. 6b**).

**Figure 6.**
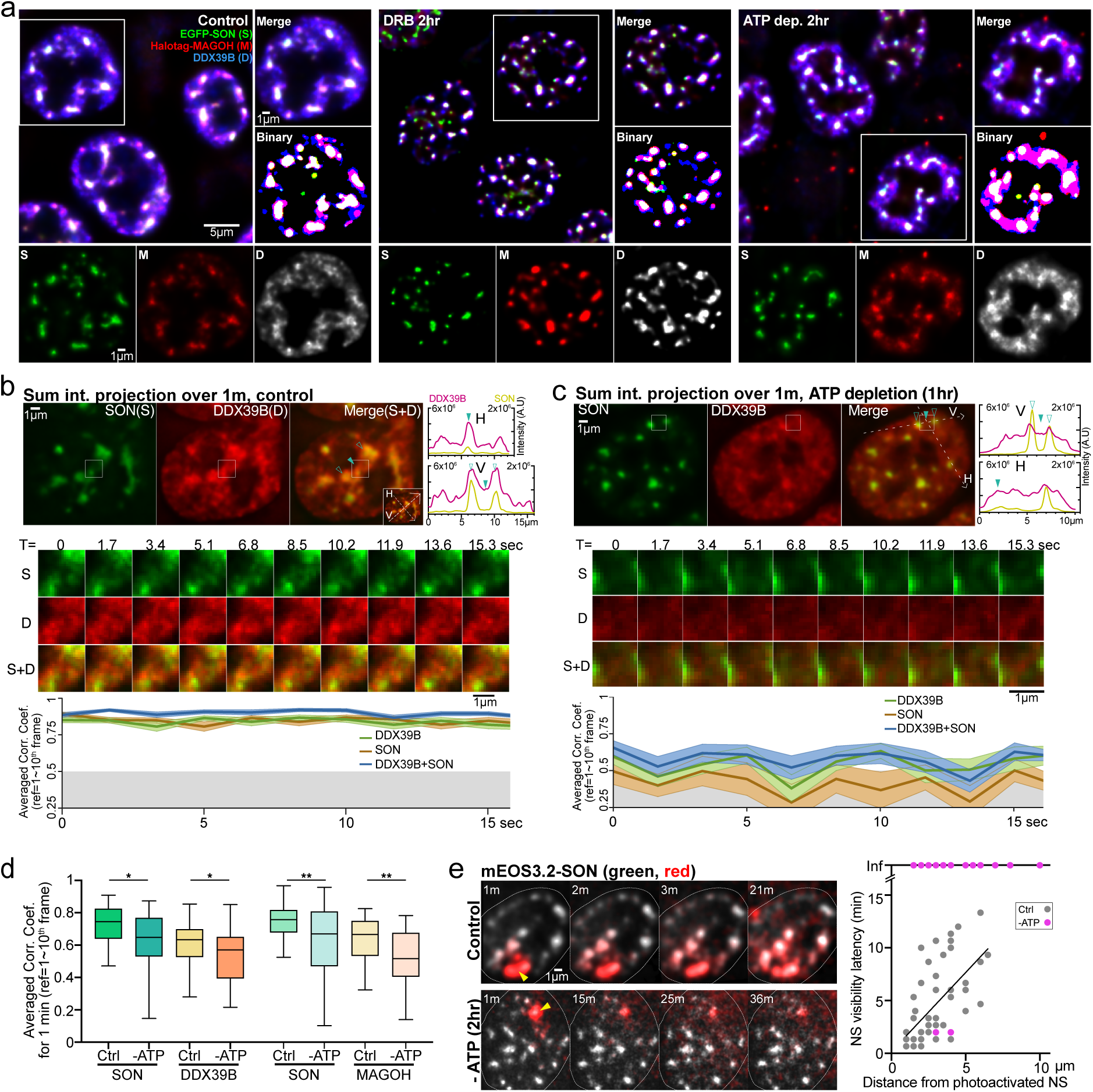
Transcription inhibition and ATP depletion perturb NS connections. **a.** Top panels: Simultaneous localization of GFP-SON (green), Halotag-MAGOH (red) and DDX39B (immunostaining) (blue) under control, 2-hr transcription inhibition by DRB, or 2-hr ATP depletion conditions. Representative images and higher-magnification views (right, enlarged insets) of the boxed nuclei showing continuous scaling (top) versus binary segmented images (bottom). Bottom panels: individual images of SON (left, green), MAGHO (middle, red), and DDX39B (right, gray) for control, DRB, and ATP-depletion conditions. Images for DRB treatment and all SON images are non-linearly scaled (*γ*=0.7). **b-c.** Dynamics of SON and DDX39B speckle connections in control (**b**) versus ATP-depleted conditions (**c**). Top: Time-integrated projection (sum-intensity) over 1 min of SON and DDX39B (left). Nonsolid arrowheads point to NS; solid arrowhead points to NS connection. Intensity profiles of SON and DDX39B were measured along the arrow crossing the nucleus and passing through NS connections either horizontally (H) or vertically (V) (right). Middle: Time-lapse images of the boxed nuclear region (white lines, top panels) show stable SON and DDX39B NS connections over time (**b**). In contrast, no stable NS connections over time are observed in the boxed region between adjacent NS in the ATP-depleted cell. Bottom: cross-correlation analysis over time, calculated from the selected region in the top panel. Mean±s.e.m calculated using 1^st^-10^th^ time frames as a reference. Gray shading shows correlation coefficient range < 0.5. (See SFig. 8 for similar MAGOH dynamics.) **d.** Averaged cross-correlation coefficients over 1 minute. N= ∼33-39 nuclear locations from ∼10-17 cells, unpaired Welch’s t-test, *p= 0.0145 (SON) 0.0492 (DDX39B), **p= 0.0074 (SON), 0.0021 (MAGOH). **e.** Loss of apparent NS connections correlates with loss of SON transfer between NS after ATP-depletion. Left: non-photoconverted mEOS3.2-SON (green, grayscale) showing NS locations versus spreading of photoconverted mEOS3.2-SON (red) from photoactivated NS (yellow arrowhead) for control (top) versus ATP-depletion (bottom) conditions. Minimal spreading of photoconverted mEOS3.2-SON was observed over 40 mins after ATP-depletion. Right: Plots of time required for photoconverted mEOS3.2-SON to reach target NS (y-axis) versus distance to target NS (x-axis) in control versus ATP-depletion conditions. "Inf" is for NS with no detectable photoconverted mEOS3.2-SON in ATP-deleted cells. N=55 NS from 16 control cells, 53 NS from 11 ATP-depleted cells. Slope= 0.2682 for NS in control cells.

Unlike their close colocalization within NS, SON and MAGOH only partially overlap within these NS connections (**Fig. 5b**, left). Overall, MAGOH is more stable over time and occupies a larger area than the regions of elevated SON intensity, which remain mostly confined within regions of elevated MAGOH intensity. This spatial distinction is better appreciated in the binary images of SON and MAGOH (**Fig. 5b**, left, 2nd from bottom panel, **SFig. 6a**). Most SON-segmented regions (green) overlap with MAGOH-segmented regions (red), creating yellow areas, though significant MAGOH-only regions also exist. While SON predominantly resides within MAGOH areas, smaller SON-only regions (green) appear within or adjacent to the MAGOH connections (e.g., T=13.4 sec). At specific time points, such as T=0 and 5.0s, contiguous yellow bridges span the NS, but in most other frames only MAGOH appears continuous, while SON is fragmented, displaying spatial gaps in the NS connection. A similar trend is observed with SON and DDX39B, with SON primarily present within broader DDX39B accumulations (**Fig. 5b**, right panel).

Despite the transient dynamics of SON, MAGOH, and DDX39B in these NS-connecting regions, they consistently formed at the same or nearby locations (**Fig. 5b-d, SFig. 6**). We assessed the recurrence of elevated protein concentrations at these locations by calculating Pearson correlation coefficients (PCC) between a reference time frame and all other time frames (**Fig. 5c**). Spatial correlation coefficients for all three proteins ranged from ∼0.5 to 0.8, with slight fluctuations on a timescale of several seconds.

Binary "summed" images (white) created by the union of segmented SON and MAGOH or segmented SON and DDX39B remained stable over tens of seconds to ∼1-3 min imaging periods (**Fig. 5b** bottom panels, **SFig. 6**). The composite structures created by summing these two proteins had higher and more constant PCC values than the PCC values of the individual proteins, averaging ∼0.7 for the SON/MAGOH and ∼0.8 for the SON/DDX39B composite images (**Fig. 5c**). As a negative control, correlation analysis of surrounding background intensities yielded correlation coefficients ranging from approximately -0.25 to 0.25, confirming the specificity of the observed spatial correlations (**SFig. 6**).

Finally, to test the degree to which the results of the correlation analysis depended on the initial choice of the reference frame, we repeated the correlation calculations using each of the first ten frames as references (**SFig. 6**). The average and overall range of correlation values using these 10 different reference frames showed little variation, further supporting our conclusions.

In summary, rather than existing as stably and spatially separated nuclear bodies, SON, MAGOH, and DDX39B formed dynamic and recurring juxtapositions that established apparent connections between NS. While each phase exhibited dynamics on a timescale of seconds, their composite connections showed significantly greater temporal stability, persisting for minutes.

### Translocations and viscoelastic dynamics of condensates within NS connections

Having described the overall dynamics of protein distributions in the NS connections, we next examined more carefully the details of these dynamics, which showed varying viscoelastic behaviors for the different multiphase components (**Fig. 5d, SFig. 7**).

**Figure 7.**
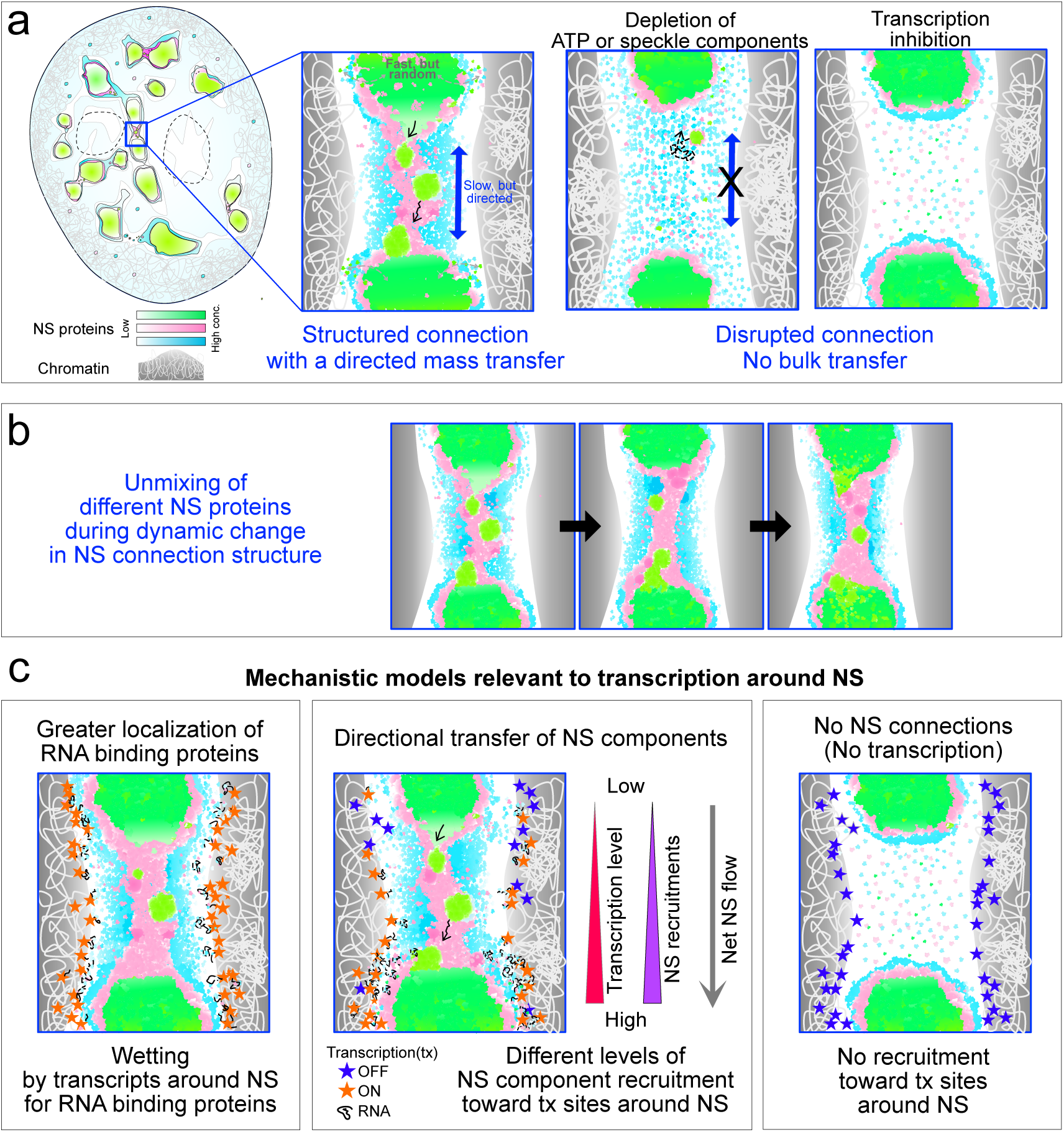
Models for NS connection dynamics and nonrandom bulk transport of NS components between adjacent NS. **a.** Multiple NS proteins form phase-separated condensates with elevated concentrations between adjacent NS within DNA-depleted regions. Fast individual protein movements within NS and NS connections coexist with slower, directional movements of condensates between NS, which are disrupted by ATP-depletion or transcriptional inhibition. **b.** Multiple NS proteins undergo unmixing within NS connections; these co-existing, but distinct condensates display correlated viscoelastic behaviors. Adhesive interactions between these condensates creates a metastable NS connection that is more stable than the individual condensates on their own. **c.** Speculative models describing how multiphase NS connections might organize to create bulk flux between NS. Left: Actively expressing genes (orange stars) position at periphery of DNA-dense regions near NS and NS connection surfaces, producing nascent RNAs (black lines) that interact (wetting) with proteins in NS and their connections. Middle: Temporal variations in net gene activity between neighboring NS create imbalances in nascent RNA levels and RNA-binding NS components. Elevated net gene activity enhances local recruitment of these components, potentially driving a capillary-like flow of NS material towards more transcriptionally active NS. Right: NS connections dissipate, and capillary flow stops without nascent RNA-mediated recruitment (wetting) of NS components.

This was most evident for SON, where small, near diffraction-limited foci translocated between NS after initially forming at the NS periphery (**Fig. 5d**, movement between NS from the top NS edge to the bottom NS edge). Over several to tens of seconds, these foci detached/dripped from the NS and translocated along NS connections, typically positioning adjacent to, surrounded by, or within MAGOH or DDX39B foci or protrusions during these translocations (**Fig. 5d, SFig. 7b**).

Whereas SON proteins typically translocated in bulk in the form of a blob, MAGOH and DDX39B formed blob-shaped foci which then elongated and dissipated within the connecting structure (**Fig. 5d, SFig. 7**). This mixed viscoelastic behavior complicated the tracking of MAGOH and DDX39B concentrations.

Throughout these dynamics, the different NS proteins did not fully mix. Instead, they maintained separate phases exhibiting distinct dynamics within the overall NS connection, as observed at WF resolution. SON and MAGOH appeared as distinct phases that came into contact and appeared to compete for space within the connecting region. For instance, SON condensates occupied regions devoid of MAGOH and vice versa (**Fig. 5b**, T=0 versus 13.4). Furthermore, the translocation of a SON blob coincided temporally with a MAGOH elongation, suggesting that the contact adhesion between the SON condensate combined with the elongating MAGOH condensate might provide the forces explaining the SON translocation (**Fig.5d,** top, T = 13.4∼16.8 s). Whereas SON and DDX39B blobs occasionally formed at the same position (**Fig. 5d**, bottom, T=13.3, 15.8 s), only SON blobs translocated along the trajectory of DDX39B. Due to the complexity of the condensate dynamics, these mass transfer events were often difficult to track. However, the overall observed patterns over time give the suggestion of bidirectional mass transfer between adjacent NS.

Overall, these findings suggest an interplay between NS proteins in establishing NS connections and facilitating the transfer of NS materials between neighboring NS. More specifically, over a timescale of seconds, NS protein condensates exhibit viscoelastic local movements including dripping, translocation, and extension between NS. We propose that the cumulative effect of these local condensate movements, each occurring over a second timescale, results in a net bulk flow of material between NS over a timescale of minutes, as observed for SON in our photoactivation experiments.

### Loss of NS connections and SON translocation between NS after transcription inhibition and ATP depletion

In the preceding sections, we showed how the coordinated dynamics of multiple NS proteins together formed multi-phase connections between adjacent NS more stable than the transient dynamics of their individual components. The movement of SON condensates within these NS connections suggested that these multiphase structures might play a role in facilitating SON movement between NS.

We therefore next asked whether the formation of these multi-phase NS connections was transcription- or ATP-dependent. First, we tested in HCT116 cells whether NS connections persisted after transcription inhibition or ATP depletion. We observed that the multiphase structure of NS connections largely disappeared upon both conditions (**Fig. 6a**).

Transcription inhibition caused SON, DDX39B, and MAGOH to concentrate more within NS, with fewer connections visualized between NS (**Fig. 6a**, middle). In live cells, the signals connecting NS were no longer present. Instead, lower-intensity, randomly distributed signals appeared without temporal correlation, resulting in the loss of distinct connecting structures outside NS in time-projected images. (**Video 6**).

ATP depletion produced a more dispersed distribution of MAGOH and DDX39B outside of NS, while SON showed an increased concentration within NS (**Fig. 6a**, right, **Video 7, SFig. 8a-d**). To estimate the change in localization of DDX39B and SON after ATP depletion, the same intensity thresholding applied for NS segmentation in control conditions was also used ATP-depletion. After ATP-depletion, the more diffuse distribution of DDX39B led to a much larger segmented area than that of SON speckles (**SFig. 8a-b**). After 120 minutes of ATP-depletion, there was a discernible decrease in the relative concentration of DDX39B within the segmented area but a statistically significant increase in SON concentration (**SFig. 8b**). Live-cell imaging confirmed these changes with ATP-depletion, showing a significant redistribution of DDX39B within the first 15 min compared to the control, followed by progressive changes from ∼15 min to 2 hrs (**SFig. 8c**). Similar changes were observed for MAGOH as seen for DDX39B (**SFig. 8d, Video 8**).

We then compared the temporal correlation of DDX39B and SON within the NS connections before and after ATP depletion. Whereas SON and DDX39B within NS connections maintained high temporal correlation under control conditions (PCC > 0.75), this temporal correlation significantly dropped after ATP depletion, even in the selected NS-connected regions showing slightly elevated signals in the ∼1minute time-integrated projection (**Fig. 6b-c**). Overall, upon ATP depletion, elevated SON signals between NS greatly diminished. In contrast, elevated DDX39B signals were spread out throughout nucleus, no longer confined to discrete structures between neighboring NS, as seen in the control. These signals were also less stable and appeared at variable locations randomly throughout the nucleus, thus producing a significant decrease in specific pattern in the time-integrated images. These random patterns further disappeared with prolonged ATP-depletion. The loss of temporal correlations for both DDX39B and MAGOH was confirmed by averaging cross-correlation coefficients across multiple observations (**Fig. 6d**).

Finally, we tested whether the loss of recurrent, structured connections between NS with ATP-depletion was accompanied by a change in the rate of bulk SON transfer between neighboring NS. We repeated the photoactivation experiments, as shown in **Fig. 3**, to visualize SON transfer between NS after ATP-depletion. Strikingly, we observed a near complete loss of photoconverted SON transfer between NS during a 40-minute imaging period (**Fig. 6e**).

In summary, transcription inhibition and ATP-depletion disrupted the temporal stability of multiphase connections between NS and loss of spatially recurrent, discrete NS connections. Additionally, ATP-depletion results in impaired bulk transfer of SON between NS within these NS connections.

## Discussion

Here we have described a novel intracellular transport phenomenon involving the micron-scale intranuclear movement of macromolecules through multiphase, highly dynamic connections between neighboring NS (**Fig. 7a**).

Our conclusion of long-range, directional bulk transport of the NS core protein SON between neighboring NS within the IC space is supported by multiple independent imaging approaches: First, WF microscopy and PALM super-resolution microscopy in live cells revealed low concentrations of SON protein bridging between neighboring NS in DNA-depleted regions. Cluster analysis of SON distribution analyzed from the PALM images further identified subsets of NS showing higher connectivity than others. Second, time-integrated images from live-cell WF microscopy confirmed the preferential connectivity, with time-lapse movies showing transient yet recurrent low-intensity accumulations of SON protein within the same DNA-depleted regions bridging specific neighboring NS.

Third, SMT of SON suggested a slow progressive translocation of SON protein between NS, distinct from the rapid, random diffusion expected for unbound SON molecules. These slow translocations occurred within regions of elevated SON intensity connecting NS, consistent with previous observations using time-lapse WF microscopy. Fourth, photoconversion experiments demonstrated a similar slow, non-diffusive, and ATP-dependent translocation of SON between NS, showing progressive spreading of photoactivated SON protein between neighboring NS over several to tens of minutes. Fifth, simultaneous two-color WF microscopy in live cells revealed the movement of small, near diffraction-limited blobs of SON protein traveling between NS. These movements occurred along and within connections between NS, which were formed by the alignment of SON, MAGOH and/or DDX39B protein condensates. Although each of these three proteins showed highly dynamic and transient distributions within these NS connections, their combined connection consisting of the union of each of these protein condensates was more stable over time than the connections formed by any individual protein.

This transport mechanism between NS is distinct from the previously described movement of small NS or NS protrusions merging with larger NS, which leads to a reduction in NS numbers upon transcription inhibition and which more closely aligns with the conventional view of biomolecular condensate dynamics involving fusion and fission events ^31^. In contrast, here we describe progressive, long-range, nonrandom transport of SON protein through specific, roughly linear connections between adjacent NS, contributing to the overall NS homeostasis throughout the nucleus. This connectivity between NS varies with the extent of net SON transfer between neighboring NS determined not only by the distance between these NS but also by the level of SON and other NS proteins involving the connectivity between these NS.

Although the physiological implications of this macromolecular transport between NS are not fully understood, we propose the following working model to suggest potential physiological mechanisms underlying this transport and to guide future studies: First, we suggest that the observed colocalization of proteins within nuclear speckles (NS), along with their partial unmixing in NS-connecting regions, can be explained by physical interactions (**Fig. 7b**). Strong adhesive interactions between proteins, interpreted as wetting, might act as the primary force retaining interacting proteins within a linear connection between NS. These physical interactions may arise from shared components or compositional similarities among NS proteins, as previously studied in Cajal bodies^40^, as well as nucleoli and stress granules^40,41^.

We propose that this wetting of adhesive macromolecules also occurs between NS components and the surface of chromatin, nascent RNA transcripts, and/or other macromolecular assemblies related to gene expression, which align along the boundaries of DNA-depleted regions containing the NS connections (**Fig. 7c**, left panel). Previous investigations have suggested that both transcription and DNA replication occur on the surface of DNA-dense regions^2,42^, while RNA polymerase II Ser2p and Ser5p foci surround NS^43-46^.

In a companion study, we show that both RNA polymerase II Ser2p foci and a subset of chromosome gene expression "hot-zones" line the periphery of these NS connections^14,39^. DDX39B is a RNA helicase involved in transcription and also a subunit within the RNA export TREX complex^47^. MAGOH is a component of the exon-junction complex (EJC)^48^, forming a heterodimer with RBM8A (Y4) that binds to RNA. We suggest that condensates of these proteins may associate with, or wet, RNA accumulations produced by actively transcribing genes.

We next propose that the wetting over these RNAs and possibly other components related to gene expression at the chromatin periphery would create a capillary force that facilitates the movement of NS components from one NS to another through the connecting structure (**Fig. 7c**, left panel). Fluctuations in the relative concentrations of RNA transcripts and other factors along the periphery of these NS connections might then drive a directional, non-equilibrium transfer of NS proteins toward a connected NS with higher net gene activity (**Fig. 7c**, middle panel). This proposed capillary flow could help steer NS components between neighboring NS and towards nuclear regions with transient increases in gene expression levels, as high nascent RNA levels are known to create or attract condensates of transcripts or splicing factors^23,49,50^. Conversely, the reduction of nascent transcripts after transcriptional inhibition would diminish the wetting of NS components over the chromatin surfaces lining these connections, thus reducing the capillary flow of NS components between NS, as observed experimentally (**Fig. 7c**, right panel).

Our working model also suggests the potential for similar bulk flow to occur between NS and active gene loci, supported by a recent study demonstrating chromatin movement driven by capillary forces generated by artificial condensates^51^.

Although we are still early in our investigation of these connections between NS, our findings already enhance our understanding of NS dynamics and intranuclear protein trafficking. We anticipate future studies exploring the physiological significance of these multiphase connections, including how the mechanisms mediating NS component transport between NS may also coordinate or regulate intranuclear RNA and gene movements. Additionally, it will be important to determine whether similar protein trafficking occurs between NS and active gene loci, potentially influencing local gene expression or RNA processing by modulating the enrichment of NS components.

## Materials and Methods

### Cell culture and cell line generation

HCT116 cells were grown in McCoy’s 5A (SH30200.FS, Cytiva) supplemented with 10% (v/v) fetal bovine serum (FBS) (F0926, Sigma-Aldrich) at 37 ℃ in a humidified incubator with 5% CO_2_. Protein tag sequences (EGFP, mEOS3.2, or Halotag) were inserted at the NH2 terminus of each endogenous coding sequence (SON, DDX39B, and MAGOH), as described previously^19^.

We selected gRNA sequences with the highest specificity and efficiency scores using CRISPR target data from the UCSC Genome Browser. gRNA DNA templates were generated by PCR using Q5 High-Fidelity DNA polymerase (M0491S, NEB) according to the GeneArt™ Precision gRNA Synthesis Kit protocol, and purified with a PCR purification kit (28104, Qiagen). The gRNA sequences used were AGAGAACGGAGCGGACGCCA for SON, CGGTTGTCTTTTGGGAGCCA for MAGOH, and TTCCCCTGCCGGCCCAGTTA for DDX39B. In vitro transcription was performed using a T7 RNA polymerase purified in the lab in the presence of a RNase inhibitor (M0307S, NEB). Template DNA was digested with DNase I (M0303S, NEB). gRNA were purified using the GeneArt™ Precision gRNA Synthesis Kit (29377, Invitrogen).

Donor DNA was synthesized by PCR from validated plasmid constructs using Q5 High-Fidelity DNA polymerase (M0491S, NEB). The Halotag-DDX39B construct was generated by replacing the EGFP sequence with the Halotag sequence in the EGFP-DDX39B construct^39^ using NheI and BspEI. Halotag-MAGOH was made by cloning MAGOH sequence with a GGGGS linker from the PPIA-mCherry-MAGOH plasmid^52^ and inserting it at the C-terminus of Halotag using EcoRI and KpnI. mEOS3.2-SON was made by replacing the YFP sequence in the YFP-SON construct^53^ with mEOS3.2 sequence cloned from the mEOS3.2-actin plasmid (57443, Addgene) using NdeI and BspEI. All final constructs were validated by sequencing and tested for proper expression and localization in HCT116 cells via transient transfection. Primers for donor DNA included ∼80 nt long overhangs with a 5’ amino nucleotide modification (amC6, IDT) to add homology arms for genomic targeting. PCR products were then gel-purified using a gel extraction kit (28704, Qiagen) to isolate PCR products of the correct size.

About 0.45 million cells were resuspended in 90 μl nucleofection buffer (VCA-1003, Lonza Nucleofector™ Kit V, Lonza) according to manufacturer’s instruction. They were mixed with 10 μl RNP solution containing 3 μg dCas9 (1081058, IDT), 2 μg sgRNA, and 5 μg donor DNA in nuclease-free water. Electroporation was performed using the Lonza AMAXA 4D-Nucleofector according to the manufacturer’s instructions. Cells were cultured for several days, then fluorescent cells were sorted using a FACS ARIA II at UIUC Biotech Center. For Halo-tag fusion proteins, cells were stained overnight in culture media with 0.1 nM JF646. Fluorescent cells were collected as a mixed clonal population, and, in some cases, then subcloned. Insertion was verified by genotyping, and localization of target proteins was confirmed by comparing with immunostaining.

Using this approach, we created the "HCT116 GFP-SON Clone C4" cell line with a biallelic EGFP tagging of SON and the "HCT116 mEOS3.2-SON Clone D4" cell line with a biallelic EOS3.2 tagging of SON. We then used this HCT116 GFP-SON Clone C4 line to create the "GFP-SON, Halotag-MAGOH" and "GFP-SON, Halotag-DDX39B" HCT116 lines corresponding to flow-sorted mixed-clonal populations of the GFP-SON C4 line with additional knock-ins of Halotag to tag MAGHO or DDX39B.

### Drug treatment

For transcriptional inhibition, cells were treated with 5,6-Dichlorobenzimidazole 1-β-D-ribofuranoside (DRB) 75ug/ml (D1916, Sigma-Aldrich or 10010302, Caymanchem) for the indicated time in growth media. DRB stock solution was prepared at 75mg/ml in DMSO. For ATP depletion, cells were treated with 5mM 2-Deoxy-D-glucose (D8375, Sigma-Aldrich) and 15mM sodium azide (71289, Sigma-Aldrich) for the indicated time in growth media. 2-Deoxy-D-glucose stock solution was prepared at 2M in PBS. Sodium azide stock solution was at 1M in PBS.

### Live cell wide-field (WF) light microscopy and image processing

Cells were seeded in 35 mm glass-bottom dishes (MatTek) or four-well Lab-Tek II Chambered Coverglass (155409PK, Thermo Fisher) and grown in culture media without phenol red for two days until they reached up to 80% confluence. For HaloTag staining, cells were incubated in growth media containing Janelia Fluor HaloTag ligands (JFX554 or JF646, Promega) at a final concentration of 0.05–50 nM for 1–24 hours. After staining with JFX554, media was replaced with dye-free media at least two hours before imaging.

We used a custom microscope equipped with a 100x/NA1.45 objective lens (UPLXAPO100XO, Olympus), a multi-wavelength LED light source (Aura Light Engine, Lumencor) equipped with 395/25, 475/28, 555/28, 634/22 nm, a motorized XY scanning stage (SCAN IM 120×80, Marzhauser), a piezo z-stage for rapid focusing (Z-Insert50, Piezoconcept), a motorized filter wheel for emission filters (X-FWR06A-E02D12, Zaber), two EMCCD cameras (iXon 897, Andor Technologies; frame-transfer mode; vertical shift speed: 0.9 μs), and a live-cell incubator chamber (H301, Okolab) maintaining humidity, temperature, and CO_2_ levels. Pixel size corresponded to 106 nm. Image collection was performed using Multi-dimensional acquisition (Micro-manager^54^). For two-color simultaneous imaging, emission signals were split at 580 nm using a dichroic beam splitter (FF580-FDi02-t3-25×36, Semrock). The separated signals were then directed to two cameras, each with a bandpass filter placed in front of the camera: either FF01-515/30-25 or FF01-525/50-25 (Semrock) on one camera and either FF01-595/31-25, (Semrock) or ET700/75m (Chroma) on the other. Two-color images of TetraSpeck beads (0.1μm, T7279, ThermoFisher) were used to translationally and rotationally align the two camera images and correct for chromatic aberration using the Huygens software.

The cell dish or Lab-Tek chamber was equilibrated on the microscope at 37°C and 5% CO2 for more than one hour. We then collected time-lapse images of multiple z-stacks and processed them using Huygens software (Scientific Volume Imaging) for deconvolution. The multiple z-stacks (z-spacing = 250 nm, ∼9-13 optical sections per z-stack) of each time frame were projected along the z-axis using maximum-intensity projection. Unless otherwise specified, all individual time-lapse images presented in figures corresponded to these maximum-intensity 2D projections of the 3D z-stacks. Time-integrated images were created by sum-intensity projection of these time-lapse images over time.

### Pseudo-color intensity mapping

Pseudo-color intensity maps were generated using MATLAB code which assigned pixel values into 10 discrete levels (https://github.com/Jiahkim/intmap.git). Typically, pixel values higher than 30-40% of the maximum intensity value in the image, which usually correspond to NS, were assigned to level 10. Pixel intensities between either zero or specified background intensity and the maximum intensity threshold were assigned to the remaining levels. The intensity map displayed the assigned levels in pseudo-color.

### Super-resolution microscopy and analysis

For single-molecule imaging, we used the WF imaging system described above, additionally equipped with a multi-line laser (405/488/561/638 nm, Cobolt Skyra, HÜBNER Photonics) for highly inclined and laminated optical sheet (HILO) illumination^37,38^, and a beam shaper (TopShape, asphericon) for flat-field illumination^55^. Z-focus was maintained using a z-drift correction module (CRISP Autofocus System, Applied Scientific Instrumentation). For 2D photoactivated localization microscopy (PALM)^56^, we acquired a single stack at one focal plane of 30,000 frames using 10–20 ms exposures; stochastic photoconversion of mEOS3.2-SON was achieved using a 405 nm LED illumination (0.1-0.5% intensity, Aura Light, Engine). Super-resolution images were reconstructed using the open-source software SMAP^57^. XY drift during data acquisition was corrected using a drift correction function in SMAP. Final localizations with precision better than 20 nm were clustered using the DBSCAN function in SMAP. Various combinations of neighborhood radius (ε) and minimum points (k) were tested.

### Photo-activation experiments

We used the WF imaging system described above, additionally equipped with a focused 405nm laser (Cobolt MLD, HÜBNER Photonics), to photoactivate a single NS. The focused 405nm beam was prepared by an additional lens with a 4f configuration. The generated beam is elongated along the z-axis (>10 um) and its diameter is ∼1 um. The lateral beam position was marked in the camera field-of-view. The target NS was positioned at this beam position using the motorized XY stage. The selected target NS in the nucleus was illuminated for 100-300 ms (50-200μW) and this was followed immediately by sequential two-color WF live-cell imaging of both photoconverted and non-converted mEOS3.2-SON. Images were processed as described above. Time-lapse images shown for photoactivation experiments are sum-intensity 2D projections of z-stacks.

### Single molecule tracking

We used the same microscope as for PALM but imaged a few well-separated molecules within regions of interest approximately the size of a cell nucleus. After a brief 405 nm LED illumination (0.1% intensity) to photoconvert mEOS3.2-SON in live HCT116 cells, we quickly assessed single-molecule spot density using 561 nm illumination (∼0.5-2 mW). We then acquired 3,000–10,000 frames of sparsely distributed single-molecules, with a 10 ms exposure time and 10.35 ms/frame imaging acquisition rate including camera readout time. To identify nuclear regions where single-molecule movements occurred, we captured WF images of mEOS3.2-SON (green) and DNA stained with SiR-Hoechst (150 nM for 2hr, SiR-DNA kit, Spirochrome) for 30-60 secs before and after single-molecule imaging. A SiR-Hoechst stock solution was prepared at 1mM in DMSO following the manufacturer’s protocol.

Single-molecule images were analyzed using an open-source single-molecule tracking software, UTrack^58^, allowing for a 2 frame gap (for red, mEOS3.2) in the appearance of individual molecules, and a neighborhood search radius of up to ∼700 nm. UTrack output tables contained x, y positions and corresponding time frame for each track. Tracks were grouped based on their localization in different nuclear regions identified by WF imaging. Diffusion coefficients for each region were then estimated separately using a custom MATLAB code. (https://github.com/Jiahkim/SMTloc.git).

More specifically, this MATLAB code includes 3 steps: 1) Segmenting nuclear regions (NS, interchromatin, DNA-compact regions) and the area outside of the nucleus based on segmenting WF time-integrated projections (sum-intensity projection)-WF images were acquired both before and after SMT. For segmentation, we selected the WF time-projection image that showed the highest spatial similarity in nuclear shape and NS patterns to the time-integrated projection of single molecule images; 2) Assigning nuclear locations for the x-y coordinates of each track and the proportion of each track that fell within the designated nuclear region-tracks located outside the nucleus were excluded from analysis; 3) Extracting track properties- to estimate diffusion coefficients, we generated a normalized histogram (probability) of frame-to-frame displacements using 20 nm bins. The histogram was then fit to a two-component diffusion model, as described elsewhere^59,60^.

For mean square displacement (MSD) analysis, we used Halotag-SON expressed from the endogenous locus. The Halotag-ligand JFX554 (10 nM, stock solution info, source) was added to media for 15 mins, followed by two 10-min washes with culture medium lacking phenol red. WF images of JFX554-SON and SiR-DNA were collected first. Single molecule images of JFX554-SON were then captured once the density of the single molecules was sufficiently low to ensure that molecular spots were spatially well-resolved after 561-nm illumination. When the density of single molecule spots was high, continuous illumination was applied to reduce it, allowing well-separated spots to be imaged. The analysis of these single molecule images was conducted as previously described, with 0 gap allowance for tracking linkage. Mean square displacement (MSD) over time was calculated and fitted to the equation MSD= 4DΔt^α 61^.

### Fixed cell immunostaining, microscopy, and analysis of NS morphology

HCT116 cells were grown on #1.5 coverslips (7229608, EMS) in 12- or 24-well plates for two days until reaching ∼80-90% confluency. Cells were fixed with 4% paraformaldehyde (PFA; P6148, Sigma-Aldrich) in PBS for 15 minutes, followed by 3x 5-minute washes with PBS. Coverslips were then mounted in Mowiol-DABCO anti-fade mount medium or processed for further immunostaining. For immunostaining, cells were permeabilized and blocked with 10% Triton X-100 (28314, ThermoFisher) and 1% BSA (37525, ThermoFisher) in PBS, followed by 3x 5-minute PBS washes. Coverslips were then incubated in primary antibody solution at an ∼1:300∼500 dilution with 1% BSA in PBS) for 18 hours at 4°C, followed by 3x 5-minute washes with PBS. Coverslips were incubated in secondary antibody solution at an 1:200∼300 dilution with 1% BSA in PBS). After 3-5x 5mintue washes, coverslips were mounted in Mowiol-DABCO anti-fade mount medium. To compare SON, MAGOH and DDX38B nuclear distribution in fixed samples, we performed immunostaining for DDX39B in HCT116 cells expressing EGF-tagged endogenous SON and Halotag-tagged endogenous MAGOH from their endogenous loci. A custom-made rabbit polyclonal primary antibody against DDX39B^39^ and Fab fragment of goat Anti-Rabbit IgG secondary antibody (111-007-008, JacksonimmunoResearch) labeled with Alexa 647 were used. For NS morphology analysis, we used the MATLAB function ‘adaptthresh,’ which applies a local adaptive thresholding algorithm to segment NS areas, and analyzed the result as described^31^.

### Image analysis for subcellular spatial correlation analysis

For correlation analysis between frames in live cell images, we selected local rectangular regions of interest (ROI) between NS, typically 8-20 pixels wide and long, depending on the length of the NS connection. Pearson correlation coefficients were calculated over time between the reference frame (0-10^th^ time frame) and other frames using the MATLAB function ‘corr2’, defined by equation (1) below where A_ij_ and B_ij_ are the pixel value at position(I, j) in time lapse images, and 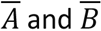 are the mean pixel value of images A and B, respectively.

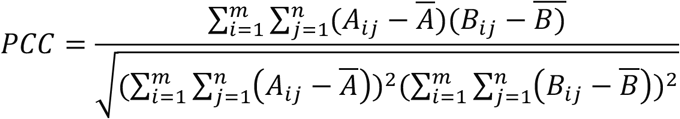

The Pearson correlation, also referred to as normalized cross-correlation, measures the linear relationship between pixel intensities in two images. Subtracting the mean and dividing by the standard deviation removes effects from differences in intensity level and contrast between the images. This normalization ensures that the PCC ranges from –1 (perfect negative correlation) to +1 (perfect positive correlation), with 0 indicating no linear correlation.

To calculate the correlation of composite structure over time, we generate the composite image by summing two rescaled target images, where pixel intensities were normalized from 0 (background) to 1 (maximum intensity within the ROI). Correlation values were calculated over time as described above.

## Supplementary Figure Legends

**Supplementary Figure 1. DBSCAN clustering analysis of SON PALM images using different parameters.**

Segmentation of molecule clusters in SON PALM images using DBSCAN showing the effects of varying two parameters. Adjusting k (minimum number of nearest neighbors) and ε (neighborhood radius, nm) affect both the size and connectivity of the detected clusters. Parameter variation leads to different levels of connectivity ranging from isolated clusters corresponding to individual NS to larger clusters corresponding to multiple NS bridged by elevated SON localization densities.

**Supplementary Figure 2. Functional variations in connectivity between neighboring NS revealed by time-integrated projection of wide-field live-cell imaging.**

Time-integrated projection (sum-intensity) over the indicated time periods ("s", secs), displayed using linear (top), nonlinear (*γ*=0.5, middle), and 10-level pseudo-color intensity mapping (bottom). Bidirectional arrows indicate NS connections, while single directional arrows indicate direction of visualized protrusions growing from one NS towards another neighboring NS. Pseudo-color scaling: 0–30% of maximum SON intensity in the field of view is divided into 10 discrete levels and color-coded as shown in the color bar. Temporal resolution is 1s per frame.

**Supplementary Figure 3. Wide-field time-lapse images of the nucleus and SON molecular movements in different nuclear regions.**

**a.** Time-lapse images (non-linearly scaled, *γ*=0.7, top) and the corresponding pseudo-color intensity mapping (bottom) of wide-field images of SON showing regions (arrows) where NS connections repeatedly form. NS connections shown in **Fig. 2b** highlighted with magenta arrows, and other detected NS connections marked with red arrows.

**b.** Randomly selected single molecule tracks (SMT) of SON from indicated time periods ("s", secs). High magnification views (bottom) of the boxed nuclear region (top). SMT are color-coded according to acquisition time (0-103 secs).

**c.** Fraction of SON molecules in the slow diffusing state (D_1_) versus fast diffusing state (D_2_) for SMT lying within nuclear speckles (NS), interchromatin space (IC), or chromatin-dense regions (C). Fractions of each state do not significantly vary in different nuclear regions. Kruskal-Wallis test (*, p<0.5).

**d.** Randomly selected SON SMT residing in the selected location (NS, IC, C) (left) or moving from one nuclear region to another, as specified (right). Top: SMT superimposed on SON or SiR-DNA WF images. Bottom: High magnification views of the boxed nuclear region (top). Few tracks move between NS and chromatin region (C).

**Supplementary Figure 4. Comparison of times for SON spreading to neighboring NS as function of straight-line versus NS connection distances**

**a.** Cartoon showing straight-line versus NS connection possible routes for photo-converted SON proteins (red) spreading from photoactivated NS (yellow lightning bolt) to adjacent NS (green) : **d**s = the straight distance regardless of chromatin barrier; **d**c = contour distance along an interchromatin channel with low density SON protein connecting adjacent NS.

**b.** Observed mean displacement distances of SON spreading between NS as a function of time for each route. ‘**d**ref ‘corresponds to cases in which there is no alternative route available (**d**s=**d**c). Linear fitting with a slope, 0.2956 for **d**ref (R^2^=0.6120), 0.1728 for **d**s (R^2^=0.2267), 0.2715 for **d**c (R^2^=0.2980).

**c.** Speed of spreading per minute calculated according to different routes. Spreading speeds calculated along straight routes show larger discrepancy from the reference speeds than those calculated along contour routes. (** *P* < 0.01 **** *P* < 0.0001; Mann-Whitney test).

**Supplementary Figure 5. Quantitative analysis of NS protein and RNA occupancy in interchromatin region identifies DDX39B and MAGOH as an extended NS marker.**

We visualized SON, SRRM2, MAGOH and DDX39B using knock-in of fluorescent tag sequence into endogenous gene loci (KI), histone modifications (H3K9ac, H3K9me3, H3K27me3), RNA pol2 S5P, and a phosphorylated epitope of SRRM2 ("SC35")^35^ using immunostaining, and MALAT1 as well as polyA-RNA using RNA FISH.

**a.** Left to right: representative images of SRRM2, MAGOH, DDX39B, H3K9ac (red, top to bottom), DNA (DAPI staining, grayscale), merged target with DNA, SON, merged target with SON, and intensity line-scans along arrows in the target protein-SON or DNA merged image.

**b.** Pearson correlation analysis between specified target image with DNA image. Bar graph: mean ± s.e.m. (n= 223, 70, 217,168, 79, 223, 79 47, 118, 200, 165 cells in the order of targets shown in **b**, respectively). Higher magnitude negative correlation coefficient indicates a higher concentration within DNA-depleted regions. A larger positive correlation coefficient indicates a greater overlap with DNA. MALAT1 and polyA-RNA signals measured by RNA FISH. KI refers to measurement of tagged proteins expressed from endogenous gene locus.

**Supplementary Figure 6. High spatial correlation between multiple proteins within NS connections in different nuclear regions**

These regions are from the boxed region in Fig. 5.

**a** & **b**. Time-lapse images and cross-correlation analysis over time for SON (S) and MAGOH (M) with a frame rate, 1.68s/frame, or DDX39B (D) with 0.83s/frame, and merge of target proteins (M+S or D+S). The line plots display mean±s.e.m calculated from 1^st^ -10^th^ time frames used as a reference. Boxplots show the mean of cross-correlation coefficient over the indicated imaging periods for SON, MAGOH or DDX39B, and merge of target proteins. “bg” is background region outside of NS connections identified by time-integrated projections. Dynamics of SON and MAGOH (region1, top panel in (**a**)) or DDX39B (region2, bottom panel in (**b**)) within NS connection show various apparent viscoelastic behaviors, including translocation of SON condensate blobs (arrowheads), dispersion of MAGOH, stretching or contraction of DDX39B (arrows) along connection.

**Supplementary Figure 7. Apparent viscoelastic behaviors of NS proteins.**

**a.** SON and MAGOH, showing protrusions/dripping, blob formation, translocation or dispersion and fusion within connections between NS. Arrowheads and circles show SON or MAGOH blobs. Right panels show binary versions of the left panels. Arrows suggest likely directions of connection formation or blob movement towards another NS.

**b.** SON and DDX39B dynamics, similar formatting as in (**a**).

**Supplementary Figure 8. ATP-depletion disperses MAGOH and DDX39B between NS with loss of metastable NS connections.**

**a.** Representative images of SON (green), DDX39B (red), DAPI (blue) in control versus ATP-depleted cells (fixed samples), and intensity line-scans measured along the white arrow in the merged image. The intensity plot is displayed using minimum and maximum intensity of DDX39B in the control cell. Gray line indicates threshold used for segmentation of DDX39B image.

**b.** Quantitative analysis of NS morphology changes between control and ATP-depleted cells. Segmented SON areas become brighter but smaller, while segmented DDX39B area become dimmer but larger. Binary images of SON and DDX39B (shown in (**a**)) were generated using the same threshold (gray line shown in plots in (**a**)) for both conditions. Green lines show the nuclear outline. Bar plots show individual segmented areas of SON and DDX39B (mean ± s.e.m.). Box plots show total intensity of segmented area or nucleoplasm (out of the segmented area) measured in ATP-depleted cells, relative to that in control cells. N > 2300 segmented area (NS) from 242 control cells, and from 237 ATP-depleted cells. ****p<0.0001, ns >0.05

**c.** Left: Time-lapse images of SON and DDX39B showing a gradual change of DDX39B morphology after ATP-deletion in live cells, relative to the morphology and dynamics observed in control cells. In contrast, NS defined by SON concentrations show noticeable shrinkage occurring within first 15 mins. Right: measured areas for segmented SON versus DDX39B regions averaged over >1000 segmented areas (NS) from 166 control cells and 137 ATP-treated cells.

**d.** Same as in (**b**), but for SON and MAGOH. N >1000 segmented areas (NS) from 135 cells. MAGOH shows progressive increase in dispersion between 0-50 mins of ATP-depletion.

**e-f.** Dynamics of SON and MAGOH speckle connections in control (**e**), and ATP-depleted cells (**f**). Top: Time-integrated projection over 1 min (sum-intensity) of SON and MAGOH. Middle: Time-lapse images of boxed regions in nuclei (top panel) showing stable NS connection dynamics in the control cell (**e**) but absence of stable connection dynamics in the ATP-depleted cell (**f**). Bottom: cross-correlation analysis over time, calculated from the same boxed regions (top panel), shows much higher average correlation coefficients for MAGOH, SON, and merged signals in the control cell (left) compared to the ATP-depleted cell (right). Mean±s.e.m calculated using 1^st^ -10^th^ time frames as a reference.

## Supplementary Information

**Video 1:** A video of endogenous EGFP-SON in live HCT116 cells with corresponding intensity mapping, showing dynamic connections between particular nuclear speckles (NS). Images were acquired at 1 s per frame.

**Video 2:** A video showing time-integrated projection of time-lapse images (1s/frame) over the indicated period, and the corresponding intensity mapping showing dynamic connectivity showing enhanced NS connections between particular NS.

**Video 3:** A video showing directional spreading of photo-converted mEOS3.2-SON, forming a connection toward a specific NS. Time-lapse images are sum-intensity projections of z-slices, displayed with a nonlinear intensity scale (*γ* = 0.6), with the corresponding intensity mapping shown on the right.

**Video 4:** A video of endogenous EGFP-SON (green), Halotag (JF646)-MAGOH (red), and the merged in live HCT116 cells. Data were acquired at 1.67 s per frame.

**Video 5:** A video of endogenous EGFP-SON (green) and Halotag (JF646)-DDX39B (red), and the merged in live HCT116 cells. Data were acquired at 0.83 s per frame.

**Video 6:** A video of endogenous EGFP-SON (green) and Halotag (JF646)-DDX39B (red) in live HCT116 cells under control condition and after transcription inhibition by > 2-hour DRB treatment. Images were acquired at 1.67 s per frame. The top two images show individual time-lapse sequences. The bottom two images show time-integrated images generated by sum-intensity projection of the corresponding time-lapse sequences shown above, the indicated time durations (+ time). All images are displayed based on its own minimum and maximum values.

**Video 7:** A video showing endogenous EGFP-SON (green) and Halotag (JF646)-DDX39B (red) in live HCT116 cells with and without ATP depletion. Images were acquired at 1.67 s per frame. The two images in the left panel are displayed using the same intensity range based on the minimum and maximum values of the control cell. The image of ATP-depleted cell on the right is displayed using a different intensity range based on its own minimum and maximum values, to visualize low intensity patterns.

**Video 8:** A video of endogenous EGFP-SON (green) and Halotag (JF646)-MAGOH (red) in live HCT116 cells with and without ATP depletion. Data were acquired at 1.67 s per frame.

## Acknowledgements

We thank U01 collaborator, Y. Shav-tal for discussion. We thank L. Lavis for providing dyes for Halotag labeling, and A.Copik for helping cell soring for cell line screening. This work was supported by U01DK127422 (ASB, KYH, YS), NIH R35GM138039 (KYH), and NIH R01GM058460 (ASB).

## Data Availability

We have provided images and live cell movies in figures and extended data figures. All the raw image data are available upon request.

## Author contribution

Conceptualization, J.K. and A.S.B.; Imaging setup, K.Y.H.; Investigation, Visualization, Analysis, J.K; Writing &Editing J.K. and A.S.B.; Cell line - SON tagging, G.H., DDX39B&MAGOH, J.K.; Initial speckle protein screening, J. K and N.C.; Funding, ASB, KYH, YS;

