## Supplementary Figures for "Nonrandom interchromatin trafficking through dynamic multiphase speckle connections"

### Supplementary Figure 1.

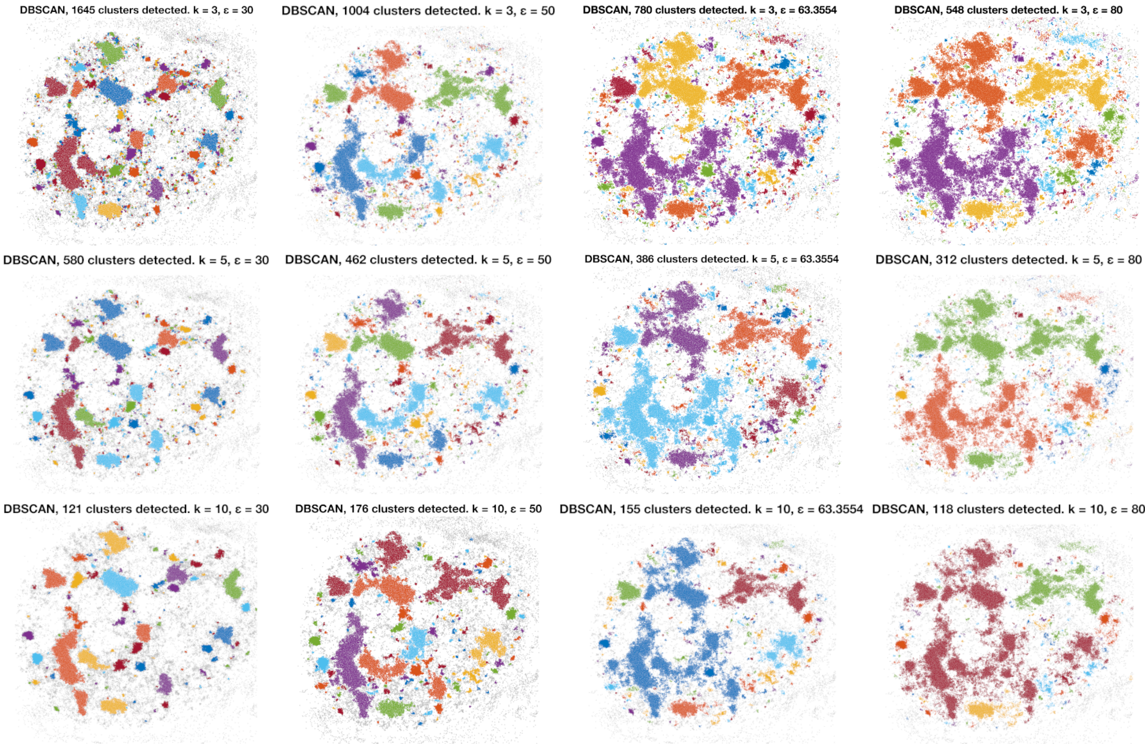

### Supplementary Figure 2

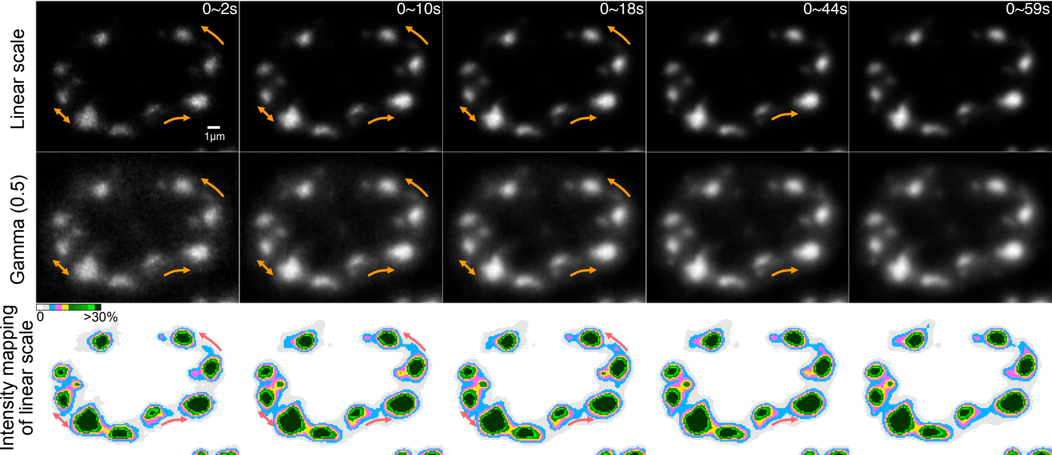

### Supplementary Figure 3

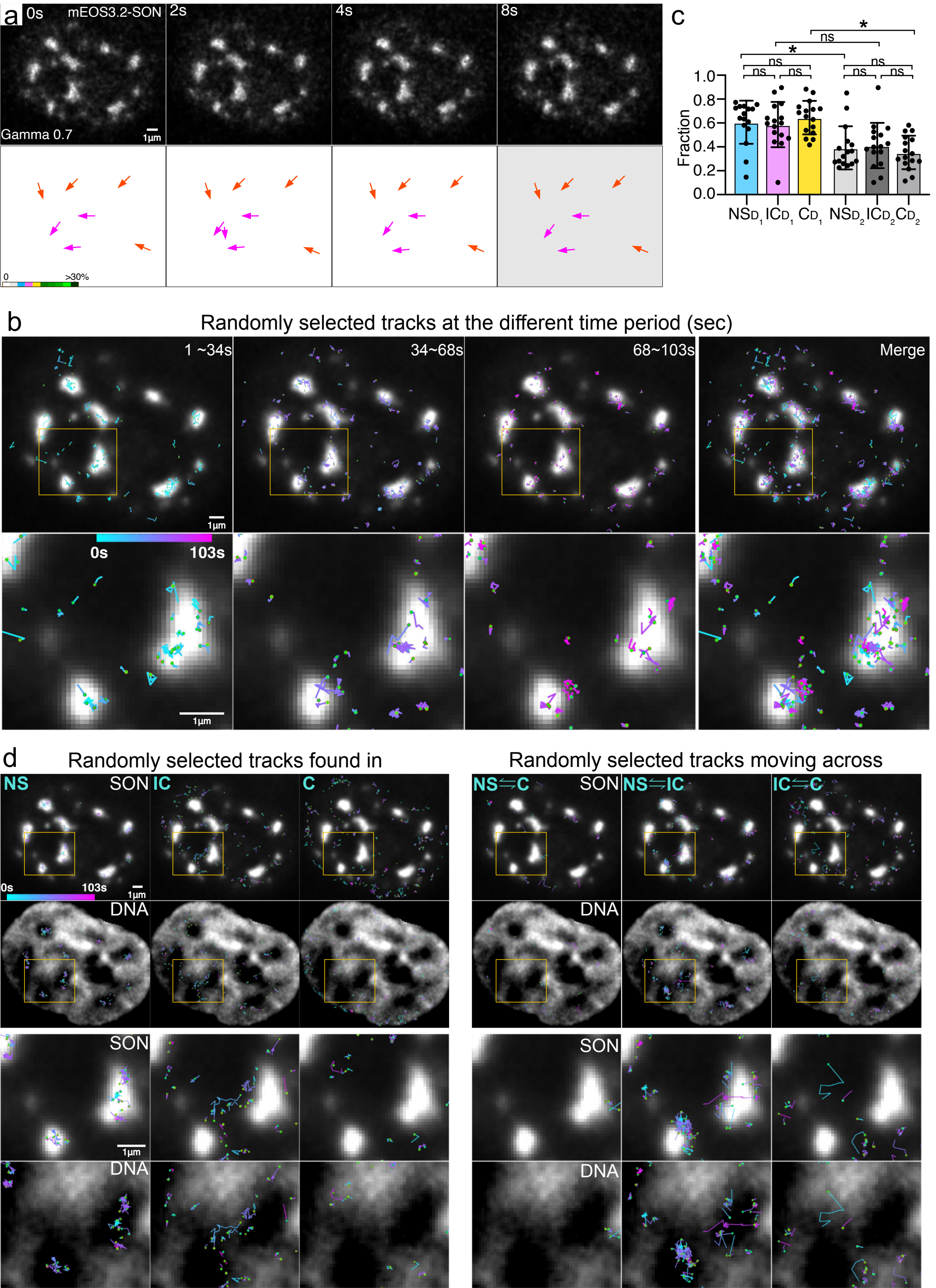

### Supplementary Figure 4

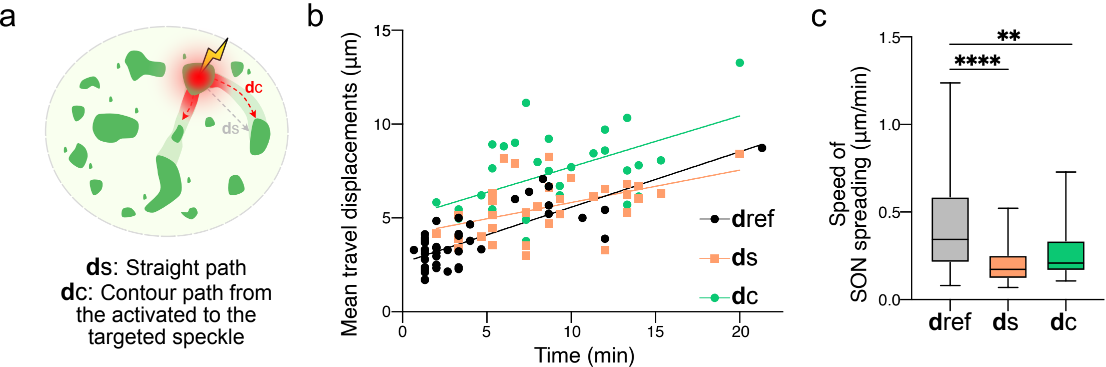

### Supplementary Figure 5

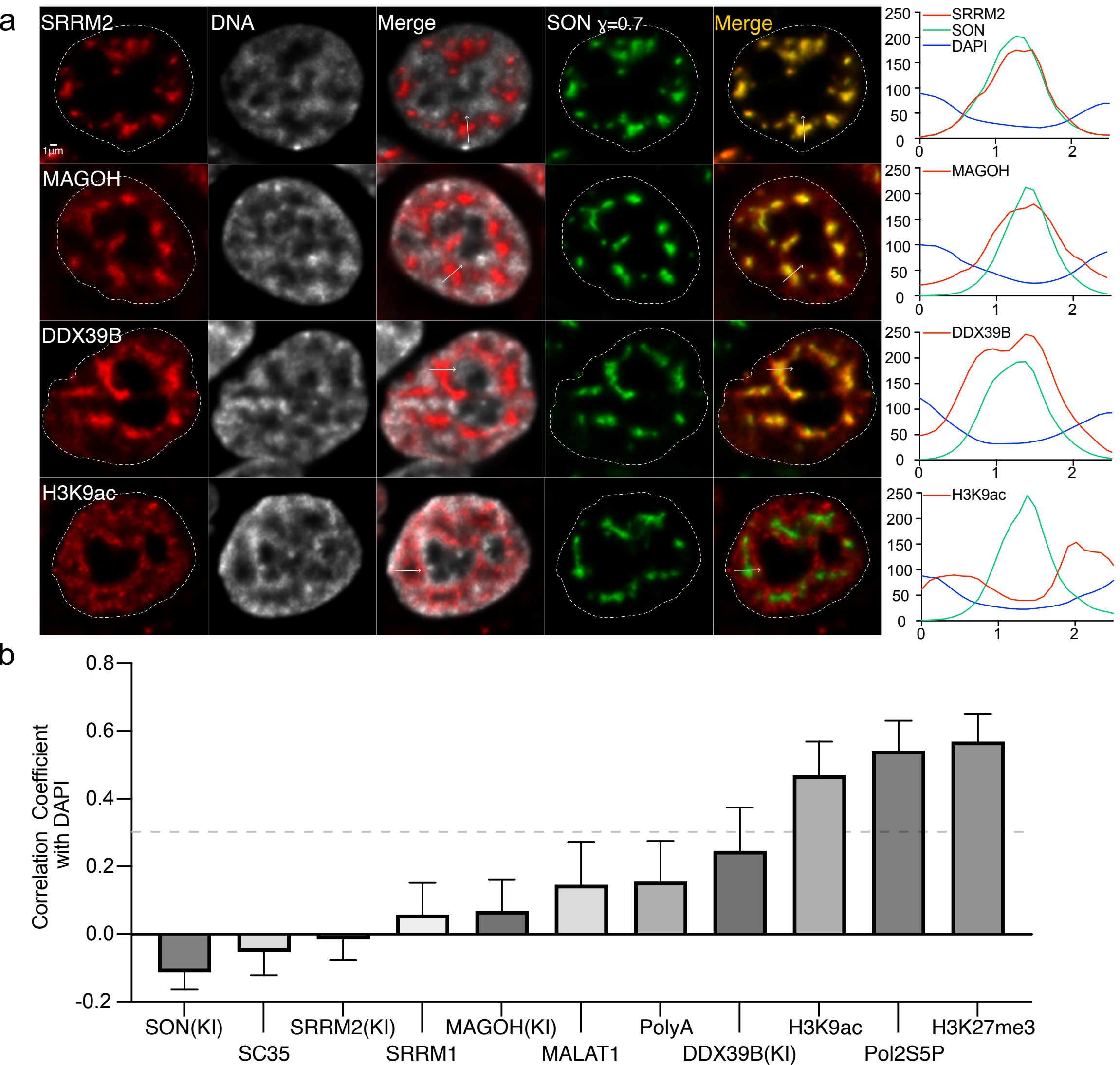

### Supplementary Figure 6

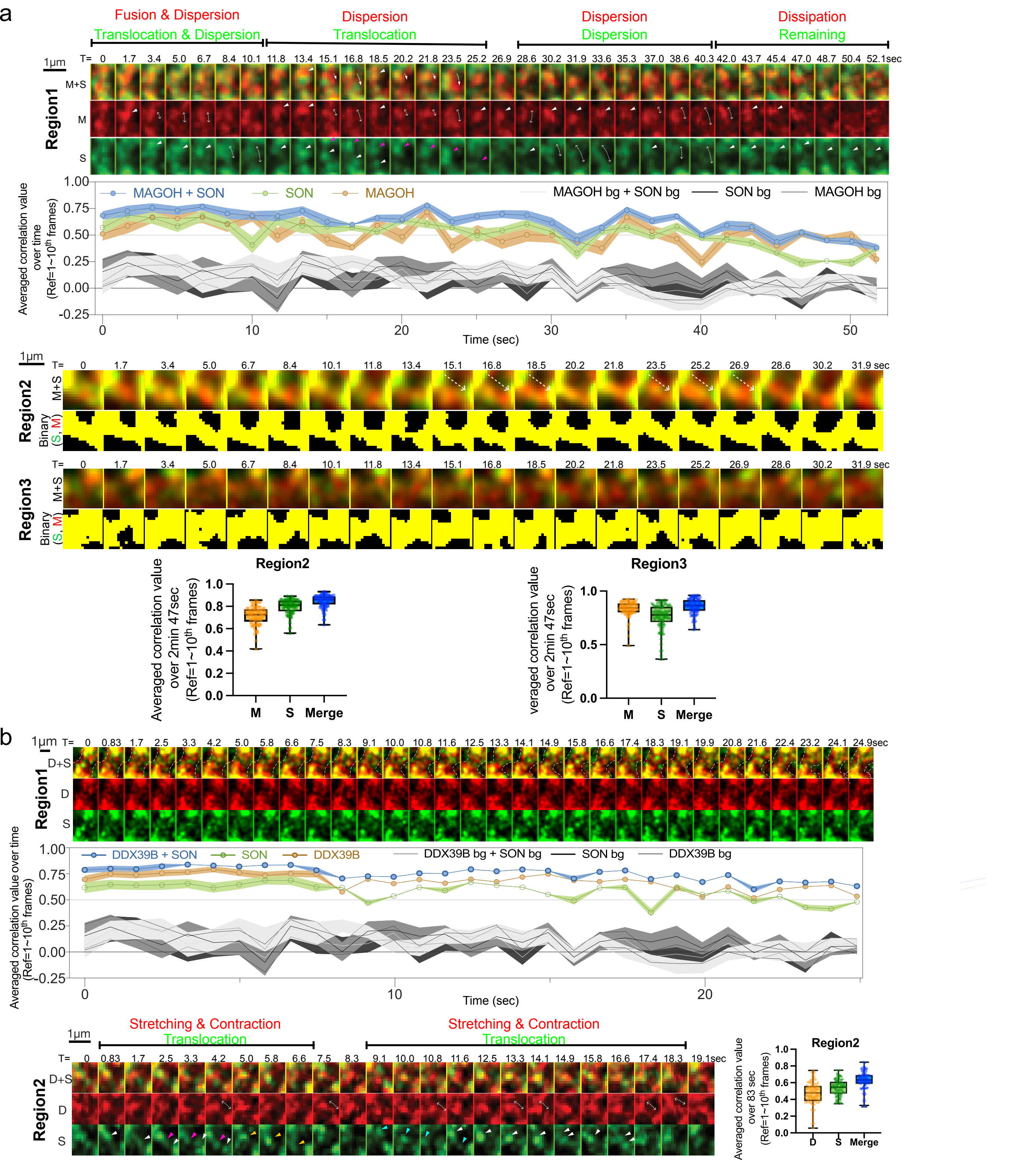

#### Supplementary Figure 7

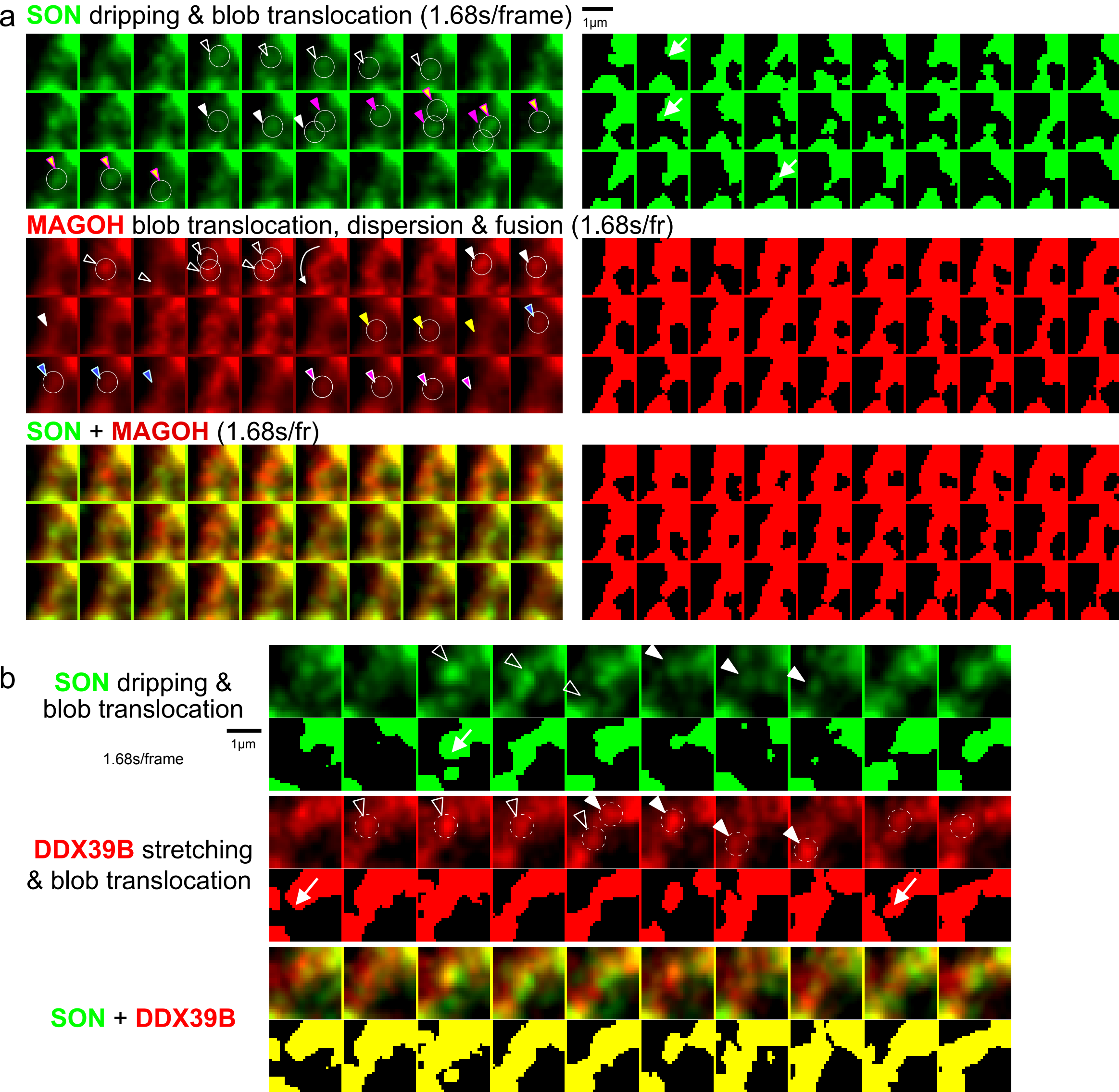

### Supplementary Figure 8

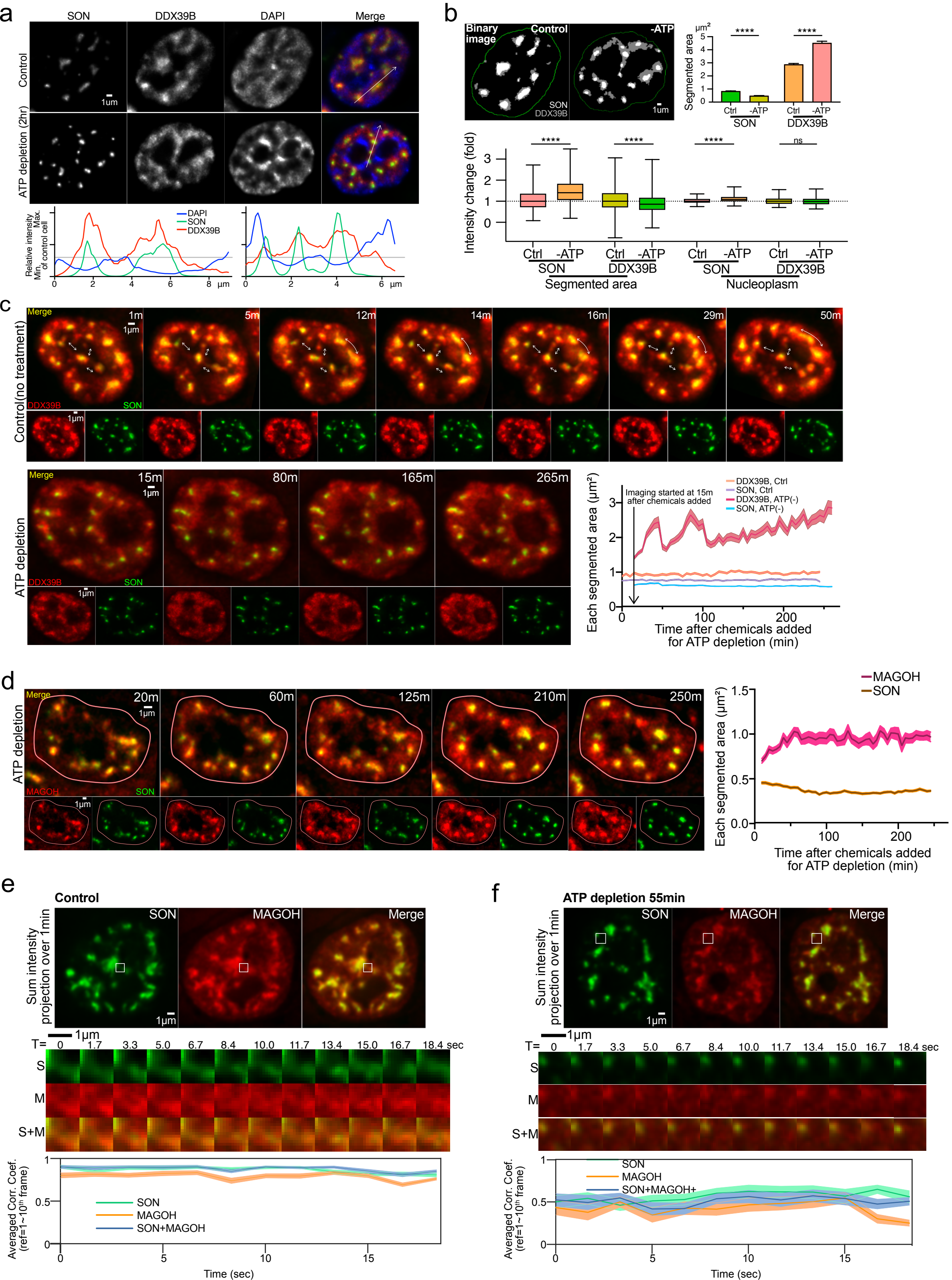
